# Dietary Restriction Enhances CD8⁺ T Cell Ketolysis to Limit Exhaustion and Boost Anti-Tumor Immunity

**DOI:** 10.1101/2024.11.14.621733

**Authors:** Brandon M. Oswald, Lisa M. DeCamp, Joseph Longo, Michael S. Dahabieh, Nicholas Bunda, Shixin Ma, McLane J Watson, Ryan D. Sheldon, Michael P. Vincent, Benjamin K. Johnson, Abigail E Ellis, Molly T. Soper-Hopper, Christine N Isaguirre, Hui Shen, Kelsey S. Williams, Peter A. Crawford, Susan Kaech, H. Josh Jang, Connie M. Krawczyk, Russell G. Jones

**Author notes:** Corresponding author: Russell G. Jones.

## Abstract

Reducing calorie intake without malnutrition limits tumor progression but the underlying mechanisms are poorly understood. Here we show that dietary restriction (DR) suppresses tumor growth by enhancing CD8^+^ T cell-mediated anti-tumor immunity. DR reshapes CD8^+^ T cell differentiation within the tumor microenvironment (TME), promoting the development of effector T cell subsets while limiting the accumulation of exhausted T (Tex) cells, and synergizes with anti-PD1 immunotherapy to restrict tumor growth. Mechanistically, DR enhances CD8^+^ T cell metabolic fitness through increased ketone body oxidation (ketolysis), which boosts mitochondrial membrane potential and fuels tricarboxylic acid (TCA) cycle-dependent pathways essential for T cell function. T cells deficient for ketolysis exhibit reduced mitochondrial function, increased exhaustion, and fail to control tumor growth under DR conditions. Our findings reveal a critical role for the immune system in mediating the anti-tumor effects of DR, highlighting nutritional modulation of CD8^+^ T cell fate in the TME as a critical determinant of anti-tumor immunity.

## Introduction

The balance between cancer cell proliferation and anti-tumor functions of the immune system influences tumor growth (*1*). Immune checkpoint inhibitors (ICIs) have revolutionized the treatment landscape for various malignancies in part by promoting the expansion of CD8^+^ T cell subsets critical for controlling tumor progression (*2*). Given their central role in mediating anti-tumor immunity, CD8^+^ T cells are often the target of immune evasion and suppression mechanisms. Chronic exposure to tumor antigens and inflammatory conditions in the tumor microenvironment (TME) can lead to CD8^+^ T cell dysfunction (also known as exhaustion), a terminally differentiated state characterized by reduced proliferative capacity and the expression of inhibitory receptors (i.e., PD1, LAG3, TIM3) that limit T cell effector function (*3*, *4*). The accumulation of exhausted T (Tex) cells in tumors is driven by the transcription factor Thymocyte Selection-Associated High Mobility Group Box (TOX) and is reinforced through epigenetic programming (*5–8*), this ultimately limits the ability of CD8⁺ T cells to control tumor growth. Understanding mechanisms that drive CD8⁺ T cell fate decisions in the TME is critical for developing new approaches to counter T cell dysfunction and overcome ICI resistance during tumor progression.

T cell dysfunction is also influenced by environmental conditions in the TME that impact T cell metabolism (*9*, *10*). CD8⁺ T cells rely on glycolysis and oxidative phosphorylation (OXPHOS) to fuel their proliferation and effector function (*11–13*). Chronic antigen stimulation, hypoxic stress, and PD1 signaling all promote metabolic and mitochondrial derangements in CD8^+^ T cells that precede their dysfunction (*14–17*). Conversely, CD8^+^ T cell effector function is enhanced by exposure to nutrients that fuel mitochondrial oxidative metabolism and acetyl-CoA levels, including ketone bodies (KBs) and acetate (*18–20*). Nutrient levels in the TME are highly influenced by diet (*21*). Thus, using dietary strategies that modulate metabolic conditions in the TME has the potential to stimulate robust anti-tumor immunity, either in isolation or in combination with ICI treatments, via metabolic enhancement of CD8^+^ T cell function.

Dietary restriction (DR), a regimen that reduces caloric intake without malnutrition, extends lifespan in mammals and delays the onset of age-related diseases including cancer (*22–26*). Dietary interventions that limit tumor growth−such as DR, fasting-mimicking diets, or low glycemic diets−are presumed to act by altering the levels of hormone (i.e., insulin, IGF-1) and/or metabolites (i.e., glucose) in the TME (*27*–*30*). However, DR also exerts major effects on immune function (*31*). Despite lower calorie intake, DR enhances both primary and memory T cell responses to bacterial infection in mice (*32*, *33*). Similarly, a 30% reduction in calorie intake is sufficient to boost the proliferation and effector function of human T cells (*34*). Yet, the contribution of the immune system to the anti-tumor effects of DR are unknown. Here, we show that DR-driven changes in ketone body (KB) metabolism reshape CD8^+^ T fate in the TME to control tumor growth.

## Results

### Dietary restriction (DR) enhances T cell-mediated anti-tumor immunity

To explore how the immune system contributes to the anti-tumor effects of DR, we examined tumor growth in immunocompetent mice using an established model of DR where daily food availability is reduced by 50% with no change to nutritional content (**Table S1**) (*33*). C57BL/6J mice were fed either *ad libitum* (AL) or subjected to 50% DR for 7 days prior to establishment of subcutaneous syngeneic tumor allografts (**Fig. 1a**). Immune profiling revealed no change in CD4^+^ T cell and NK cell populations but increased numbers of CD8^+^ T cells, eosinophils, and neutrophils in the spleen of DR-conditioned mice compared to AL-fed controls after 1 week on the diet (**Table S2**). Mice on the DR regimen displayed normal levels of physical activity (**Fig. S1a**) but consumed 50% less food and water on average (**Fig. 1b**, **S1b**), resulting in a ∼15-20% loss in body weight compared to AL-fed animals (**Fig. 1c**, **S1c**). Despite this weight loss, syngeneic melanoma (B16) and breast cancer (E0771) tumors grew more slowly in mice adapted to DR, corresponding to a 30-80% extension in tumor-free survival (**Fig. 1d-e**). These results from DR are comparable to the delay in tumor growth observed in animals fed a fasting-mimicking diet (FMD) (*35*). To interrogate the involvement of the immune system in the anti-tumor effects of DR, we first assessed B16 melanoma tumor growth in *Rag2*^-/-^ mice that lack mature T and B cells. Strikingly, the delay in tumor growth promoted by DR was lost in *Rag2*-deficient animals (**Fig. 1f**). Similarly, there was no difference in B16 tumor growth in athymic nude mice (that lack mature T cells) fed AL versus the DR diet (**Fig. S2a**). Finally, depleting CD8^+^ T cells accelerated B16 tumor growth under AL conditions but also eliminated the tumor-delaying effects of DR (**Fig. 1g, S2b**). Together, these data establish that DR limits tumor growth by promoting anti-tumor CD8^+^ T cell responses.

**Figure. 1.**
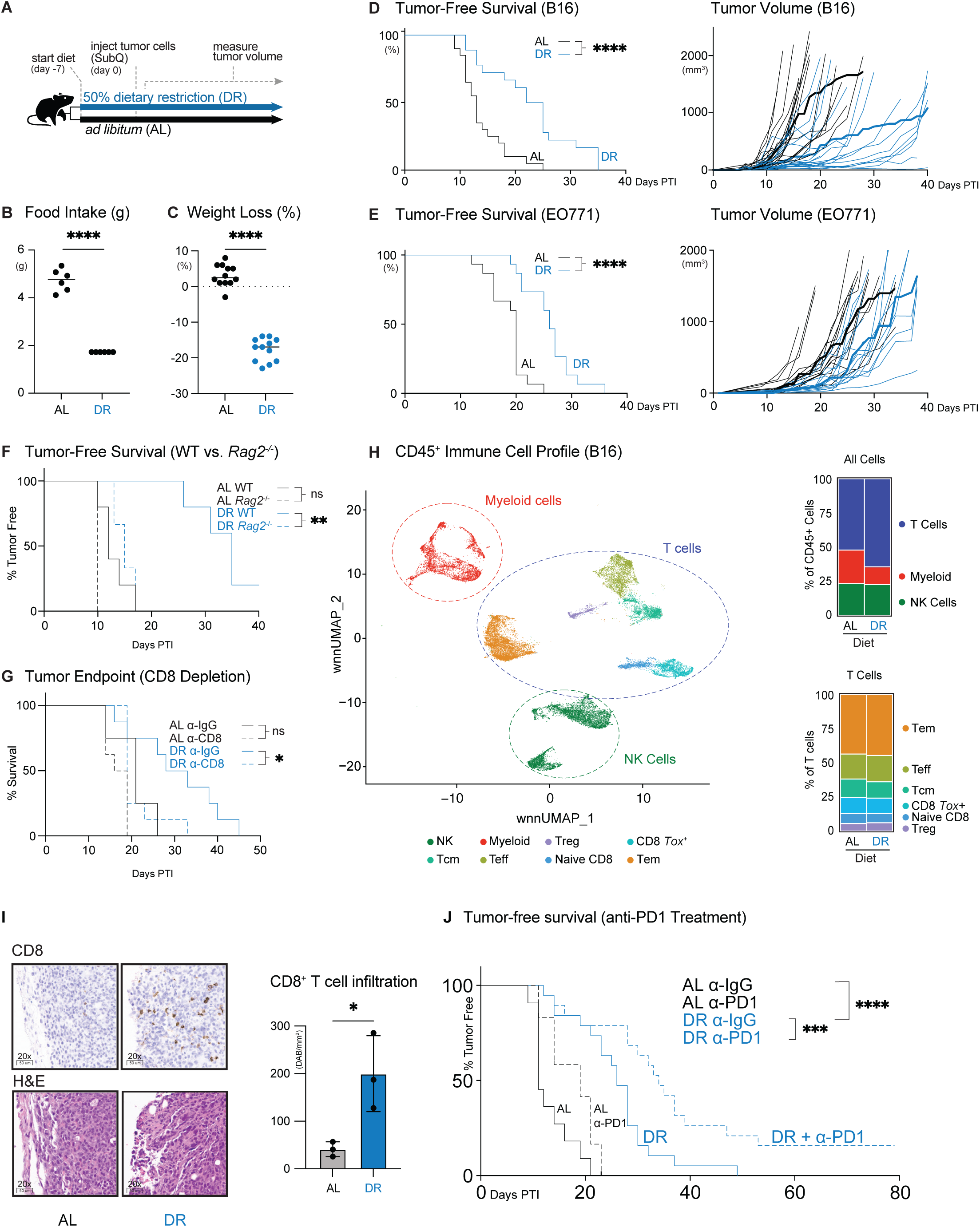
Dietary restriction enhances T cell-mediated anti-tumor immunity. A) Schematic representation of dietary interventions and tumor experiment design. B) Daily food intake (grams) of mice on *ad libitum* (AL) or dietary restriction (DR) feeding regimens. Data represent the mean ± SEM (n = 6 mice/group). C) Weight loss (% change from baseline) of mice after 1 week of feeding on AL or DR diets. Data represents the mean ± SEM (2 weeks, n = 12 mice/group). D-E) Growth of (D) B16 melanoma or (E) EO771 breast cancer cells in C57BL/6 mice fed an AL or DR diet. *Left,* Kaplan-Meier plots comparing tumor onset (tumor volume ≥250 mm³) between AL and DR groups. *Right*, spaghetti plots of individual tumor volumes growth curves over time. The darker lines indicate the average tumor growth for each diet group over time post tumor injection (PTI). B16, n=16-20 mice/group; EO771, n=15 mice/group. Statistical significance was assessed by long-rank test. F) Kaplan-Meier plot comparing tumor onset in B16 tumor-bearing wild-type (WT) and *Rag2*^-/-^ mice on an AL or DR diet. Statistical significance was assessed by log-rank test (n=5 mice/group). G) Kaplan-Meier plot comparing time-to-humane endpoint in CD8^+^ T cell-depleted mice fed an AL and DR diet. Mice were treated with control (IgG) or CD8⁺ T cell depleting (Anti-CD8) antibodies prior to tumor cell implantation. Statistical significance was assessed by log-rank test (n = 4-8 mice/group, endpoint = 1500mm^3^). H) Weighted nearest neighbor Uniform Manifold Approximation and Projection (wnnUMAP) of 45,455 CD45⁺ tumor-infiltrating cells (AL and DR combined) from B16 melanoma tumors (*left*; n = 4 mice/diet). *Right,* Breakdown of immune cell populations from all CD45^+^ cells (top) or CD3^+^ T cells (bottom). I) Histology of B16 tumors from AL- or DR-fed 14 days post tumor implantation. *Left,* immunohistochemical staining for CD8⁺ T cells and H&E staining of representative tumor sections. *Right,* quantification of positive CD8^+^ staining per tumor section (DAB/mm^2^). Data represent the mean ± SEM (n = 3 mice/group). J) Kaplan-Meier plot comparing B16 melanoma tumor onset in AL- or DR-fed mice that received anti-PD1 or IgG control antibodies by i.p. injection (200 μg/dose). Antibody treatment was administered every 3 days for a total of 5 injections, beginning 7 days PTI (n = 11-20 mice/group). Statistical significance was assessed by log-rank test with Bonferroni correction. *P<0.05, **P<0.01, ***P<0.001, ****P<0.0001.

To determine how DR impacts immune cells in the TME, we performed cellular indexing of transcriptomes and epitopes coupled to next generation sequencing (CITE-seq) on live CD45^+^ cells sorted from B16 tumors from AL- or DR-fed mice. This analysis identified several cell clusters that categorized into three main groups: myeloid cells, T cells, and NK cells (**Fig. 1h, S3a-b; Table S3**). The proportion of tumor-infiltrating T cells (of CD45^+^ cells) increased under DR, marked by an increase in *Gzmb*^+^ CD8^+^ T effector (Teff) cells (**Fig 1h, S3c**). Histological analysis confirmed this increase in CD8^+^ T cell infiltration into B16 tumors under DR conditions (**Fig. 1i, S3d**). The proportion of myeloid cells in B16 tumors also changed in DR-fed mice, marked by a decrease in the proportion of immunosuppressive *C1q*^+^ tumor-associated macrophages (TAM) and an expansion of *Ly6C*^+^ inflammatory monocytes (**Fig 1h, S3e**). While recent work suggests that DR can boost natural killer (NK) cell-mediated anti-tumor immunity (*36*), tumor-infiltrating NK cell populations were similar between dietary conditions (**Fig. 1h**).

Overcoming resistance to immune checkpoint inhibitors (ICIs) is a major challenge for cancer treatment. Given that DR enhances T cell-mediated control of tumor growth, we tested the efficacy of combining DR with anti-PD1 immunotherapy in mice bearing B16 melanoma tumors, which display resistance to PD1 blockade (*37*). *Ad libitum* or DR-fed C57BL/6 mice were administered anti-PD1 or IgG control antibodies beginning once tumors were palpable (∼7 days post implantation). DR synergized with anti-PD1 treatment, significantly extending the tumor-free survival of animals over anti-PD1 treatment (under AL feeding) or DR treatment alone (**Fig. 1j**). Strikingly, ∼15% of animals treated with DR plus anti-PD1 immunotherapy remained tumor-free after 80 days (**Fig. 1j**). Thus, DR enhances the efficacy of ICI treatment even in tumors refractory to conventional immunotherapies.

### Dietary restriction reduces T cell dysfunction in the tumor microenvironment

Our data point to DR affecting CD8^+^ T cell populations in the tumor microenvironment (TME). Thus, we used our CITE-seq datasets to characterize how DR impacts CD8^+^ tumor infiltrating lymphocyte (TIL) populations. Cell clusters were classified based on gene features, gene density, antibody-derived tags (ADTs), and MSigDB pathway analysis (**Fig. S4a**). Cellular indexing of both protein epitopes and transcriptional profiling revealed two major CD8^+^ TIL populations in B16 tumors based on gene expression for *T*ox, a key transcriptional regulator of T cell exhaustion (*5*, *6*) (**Fig. 2a**). The *Tox*^Low^ population consisted of five main subclusters including two *Mki67*^+^ proliferating Teff cell populations (Prolif1 and Prolif2), two progenitor exhausted T cell populations (Tex^Prog1^, Tex^Prog2^), an intermediate exhausted (Tex^Int^) population, and a small naïve-like cell population (**Fig. 2a**, **Fig. S4a-c**). *Tox*^High^ CD8^+^ T cells displayed elevated transcript levels for inhibitory receptor genes (*Lag3*^+^*Havcr2*^+^*Pcdc1*^+^), but clustered into two distinct groups: a conventional terminally exhausted (Tex^Term^) population and a proliferating (*Mki67*^+^) effector-like exhausted population (Tex^Eff^) (**Fig. 2a-b**, **Fig. S4a-c**) (*3*). <u>F</u>ast <u>G</u>ene <u>S</u>et <u>E</u>nrichment <u>A</u>nalysis (FGSEA) identified the Tex^Eff^ cluster as having the most effector-like properties, including enrichment for immune response and cell cycle genes (**Fig. 2b**). Tex^Eff^ cells also displayed increased expression of genes encoding effector molecules (i.e., *Ifng*, *Tnfa*, and *Gzmb*) (**Fig. 2b**), despite elevated expression of transcripts encoding *Tox* and inhibitory receptors (i.e., *Pdcd1*, *Lag3*, *Havcr2*) (**Fig. S4a**).

**Fig. 2.**
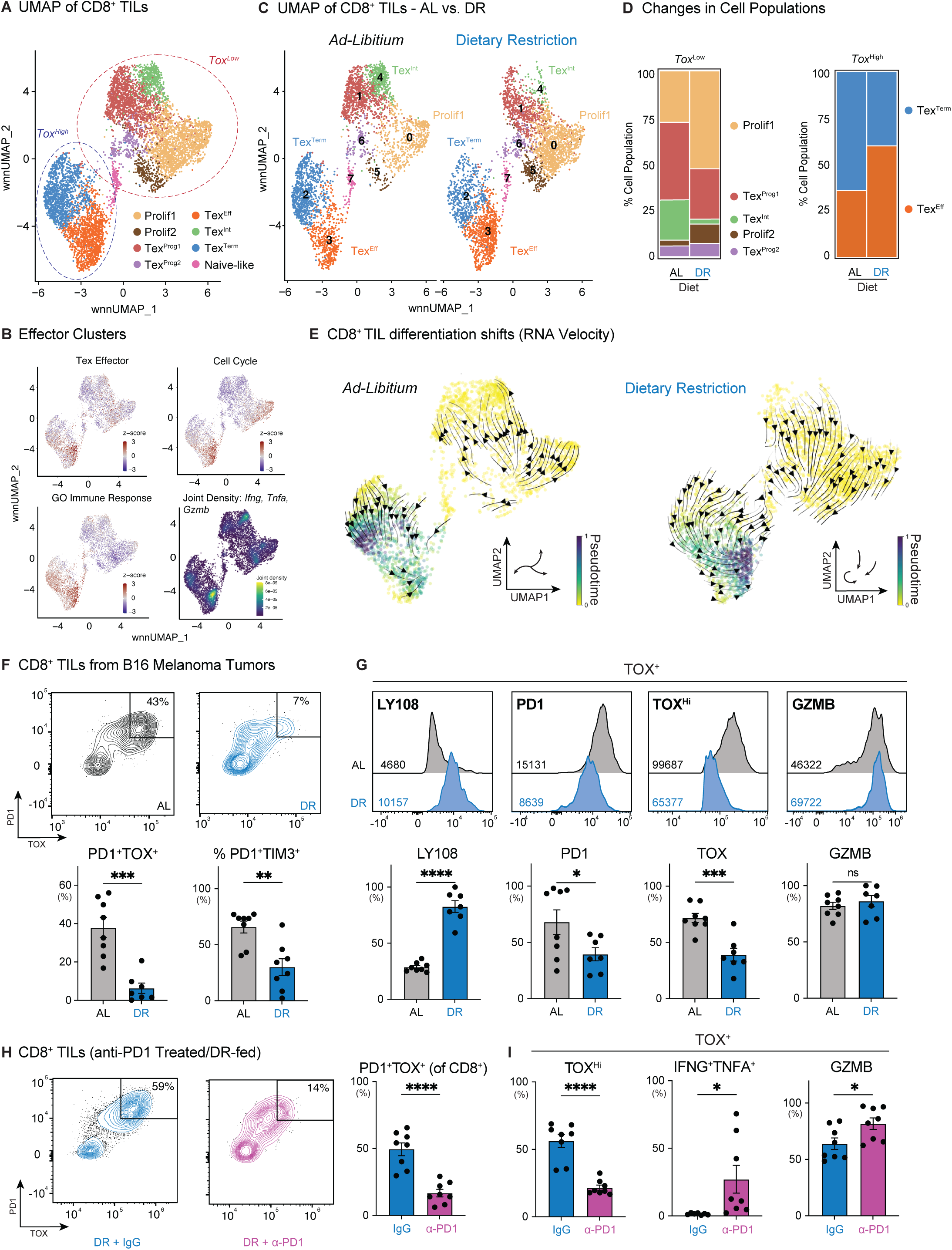
Dietary restriction reduces T cell dysfunction in the tumor microenvironment. A) Weighted nearest neighbor UMAP (wnnUMAP) of 7,005 activated (CD44^+^) CD8⁺ tumor-infiltrating lymphocytes (TILs) from B16 melanoma tumors (combined AL and DR) (n = 4). Legend indicates unique T cell clusters called based on RNA expression and antibody-derived tags (ADTs). T cell clusters with high and low *Tox* expression are highlighted by circles. B) Overlay of wnnUMAP for activated CD8^+^ TIL from (A) and MSigDB gene expression signatures for effector-like Tex cells (Tex^Eff^), Cell Cycle, and GO Immune Response pathways for TILs isolated from B16 tumors (combined AL and DR). Joint density plot indicates the highest combined expression/cell of *Ifng*, *Tnfa*, and *Gzmb* among activated CD8⁺ T cells from B16 tumors. C) wnnUMAP of activated CD8⁺ TILs divided by dietary conditions (AL: 3,348 cells, DR: 3,657 cells). Prominent CD8⁺ T cell clusters are indicated. D) Stacked bar graphs showing the percentage of CD8⁺ T cell populations divided by *Tox* expression (*Tox*^Low^ versus *Tox*^High^). E) RNA velocity plots inferring cellular differentiation trajectory for CD8⁺ T cells infiltrating B16 tumors from AL- or DR-fed mice. Trajectories were derived from the dynamical prediction model scVelo, with representative directionality for T cells under each diet shown in inset. F) PD1 and TOX expression in B16 tumor-infiltrating CD8⁺ T cells isolated from AL or DR mice 14 days PTI. *Top*, representative flow cytometry plots for PD1 versus TOX expression. *Bottom*, bar graphs showing the percentage of PD1⁺TOX⁺ and PD1⁺TIM3⁺ CD8⁺ T cells isolated from tumors. Data represents the mean ± SEM (n = 8 mice/group). G) Expression of stemness, exhaustion, and effector molecules in TOX⁺ CD8⁺ T cells isolated from B16 tumors from AL- or DR-fed mice. *Top*, representative histograms of LY108, PD1, TOX, and GZMB expression. Geometric mean fluorescence intensity (gMFI) averaged across all biological replicates is indicated in inset. *Bottom*, bar graphs quantifying the percentage of CD8^+^ cells expressing each of the indicated proteins. Data represent the mean ± SEM (n = 7-8 mice/group). H) PD1 and TOX expression in B16 tumor-infiltrating CD8⁺ T cells from DR-fed mice treated with anti-PD1 immunotherapy. DR-fed mice harboring B16 tumors were administered anti-IgG or anti-PD1 antibodies (200 ug i.p.) every 3 days for 5 doses, beginning on day 7 PTI. CD8⁺ T cells were isolated from B16 melanoma tumors at 21 days PTI. *Left*, representative flow cytometry plots for TOX versus PD1 expression. *Right*, bar graph showing the percentage of PD1⁺TOX⁺ CD8⁺ T cells following IgG or anti-PD1 treatment. Data represent the mean ± SEM (n = 8 mice/group. I) Bar graphs quantifying the percentage of TOX^+^ CD8⁺ T cells expressing high levels of TOX (TOX^High^), both IFN-γ and TNF-α (IFNG⁺TNFA⁺), and GZMB following IgG or anti-PD1 treatment from (H). Data represent the mean ± SEM (n = 8 mice/group). *P<0.05, **P<0.01, ***P<0.001, ****P<0.0001.

Single cell hashtagging strategies allowed us to track CD8^+^ TIL populations in B16 tumors from either AL- or DR-fed mice. Analysis of CD8^+^ TIL populations revealed two major trends induced by DR feeding. First, for the *Tox*^Low^ CD8^+^ TIL subset we observed an expansion in proliferating effector T cell populations (Prolif1, Prolif2) and a decrease in Tex^Int^ cells in animals conditioned to DR (**Fig. 2c-d**). Second, the type of *Tox*^High^ Tex cells found in tumors was highly dependent on diet. Tumors from AL-fed mice contained more terminally exhausted (Tex^Term^) CD8^+^ T cells, while tumors from DR mice contained a greater proportion of effector-like cells (Tex^Eff^) (**Fig. 2c-d**). To determine whether the diet-induced changes in Tex cell populations reflect differences in CD8^+^ T cell differentiation, we employed RNA velocity analysis, a computational method that infers cellular differentiation trajectories based on the abundance of spliced and unspliced mRNA in single cells (*38*). Under AL conditions, RNA velocity pseudotime (overlaid with velocity vectors) showed a directional flow away from the proliferating effector population towards Tex^Int^ and ultimately Tex^Term^ populations (**Fig. 2e**). In contrast, under DR the direction of CD8^+^ T cell differentiation culminated in the effector-like Tex^Eff^ population (**Fig. 2e**). This change in differentiation trajectory corresponded to a major shift in the ratio of effector to terminally exhausted T cell populations in B16 tumors, moving from an equal effector to terminally exhausted ratio under AL-fed conditions to a 4:1 ratio under DR (**Fig. S5a**). Collectively, these results indicate that DR triggers a major shift in CD8^+^ TIL fate away from terminal exhaustion towards a more effector-like state that favors tumor control.

We next used flow cytometry to characterize the impact of DR on CD8 T cell function within the TME. Consistent with our CITE-seq analysis, the number of CD8^+^ Tex^Term^ cells−as determined by reduced TOX and inhibitory receptor (PD1^+^TIM3^+^) expression−decreased significantly in tumors from DR-conditioned animals compared to AL-fed mice 14 days post-tumor implantation (**Fig. 2f**). Using conventional markers for terminally-exhausted (LY108^-^TIM3^+^) versus progenitor exhausted (LY108^+^TIM3^-^) cells (*2*), the majority of PD1^+^ CD8 TIL in B16 tumors from DR-fed mice were progenitor exhausted or an intermediate phenotype (LY108^+^TIM3^+^) (**Fig. S5b**) (*2*). In addition, the percentage of effector CD8^+^ T cells (PD1^low^TOX^low^) was significantly increased under DR (**Fig. 2f, S5c**) and these cells displayed increased features of stemness (LY108/SLAMF6 expression, **Fig. S5d**) and cytotoxic function (Granzyme B production, **Fig. S5e**). TOX-expressing CD8^+^ TILs from DR-fed mice displayed increased expression of stemness markers (LY108/SLAMF6) and effector molecules (Granzyme B), while showing reduced features of terminal exhaustion (i.e., lower PD1 and TOX expression) compared to AL-fed controls (**Fig. 2g**). Thus, DR both enhances the function and number of effector CD8^+^ T cells in the TME and promotes the accumulation of TOX^+^ CD8^+^ T cells that are more stem-like, less exhausted, and functionally active.

Proliferative Tex cells expressing LY108/SLAMF6 are highly responsive to ICI therapy (*2*, *39*, *40*). Given that DR synergizes with anti-PD1 treatment to slow tumor growth (**Fig. 1j**), we investigated the impact of DR on anti-tumor CD8^+^ T cell populations following ICI treatment. B16 tumor-bearing C57BL/6J mice subjected to DR were administered either anti-PD1 or control IgG antibodies 7 days post-tumor implantation, and CD8^+^ TIL phenotypes analyzed after 14 days of treatment (**Fig. S6**). Anti-PD1 ICI therapy resulted in a ∼3-fold decrease in TOX^+^PD1^+^ CD8^+^ TIL frequency (**Fig. 2h**). Moreover, TOX-expressing CD8^+^ TILs were more functional following anti-PD1 treatment, marked by lower TOX expression and increased polyfunctionality (i.e., IFN-γ^+^TNF-α^+^ and Granzyme B^+^ T cells) (**Fig. 2i**). Thus, anti-PD1 treatment amplifies the effects of dietary restriction on CD8^+^ T cell fate within the TME, favoring the expansion of CD8^+^ Teff and Tex^Eff^ cells.

### Dietary restriction enhances CD8^+^ T cell metabolic fitness via ketone body metabolism

CD8^+^ T cells are capable of metabolizing a diverse set of metabolic substrates to fuel their growth and function (*18*, *41*, *42*). Given the impact of DR on CD8^+^ T cell fate in the TME, we assessed whether DR alters the metabolic programming of CD8^+^ TIL subsets that are important for tumor control. We first conducted metabolism-focused Gene Set Enrichment Analysis (GSEA) on our CITE-seq datasets to define the metabolic features of CD8^+^ TIL subsets. This analysis revealed two distinct metabolically active CD8^+^ TIL populations in B16 tumors. CD8^+^ T cells with effector properties (Prolif1, Tex^Eff^) displayed the highest glycolytic signature (**Fig. 3a**), consistent with their high proliferative rate (**Fig. 2a-b**). Notably, Tex^Eff^ cells−which preferentially expand in tumors under DR (**Fig. 2c-d**)−displayed the highest OXPHOS signature of all CD8^+^ TIL subsets (**Fig. 3a**). Together, these results indicate that the most metabolically active CD8^+^ TIL subsets are the effector populations enhanced by DR treatment.

**Fig. 3.**
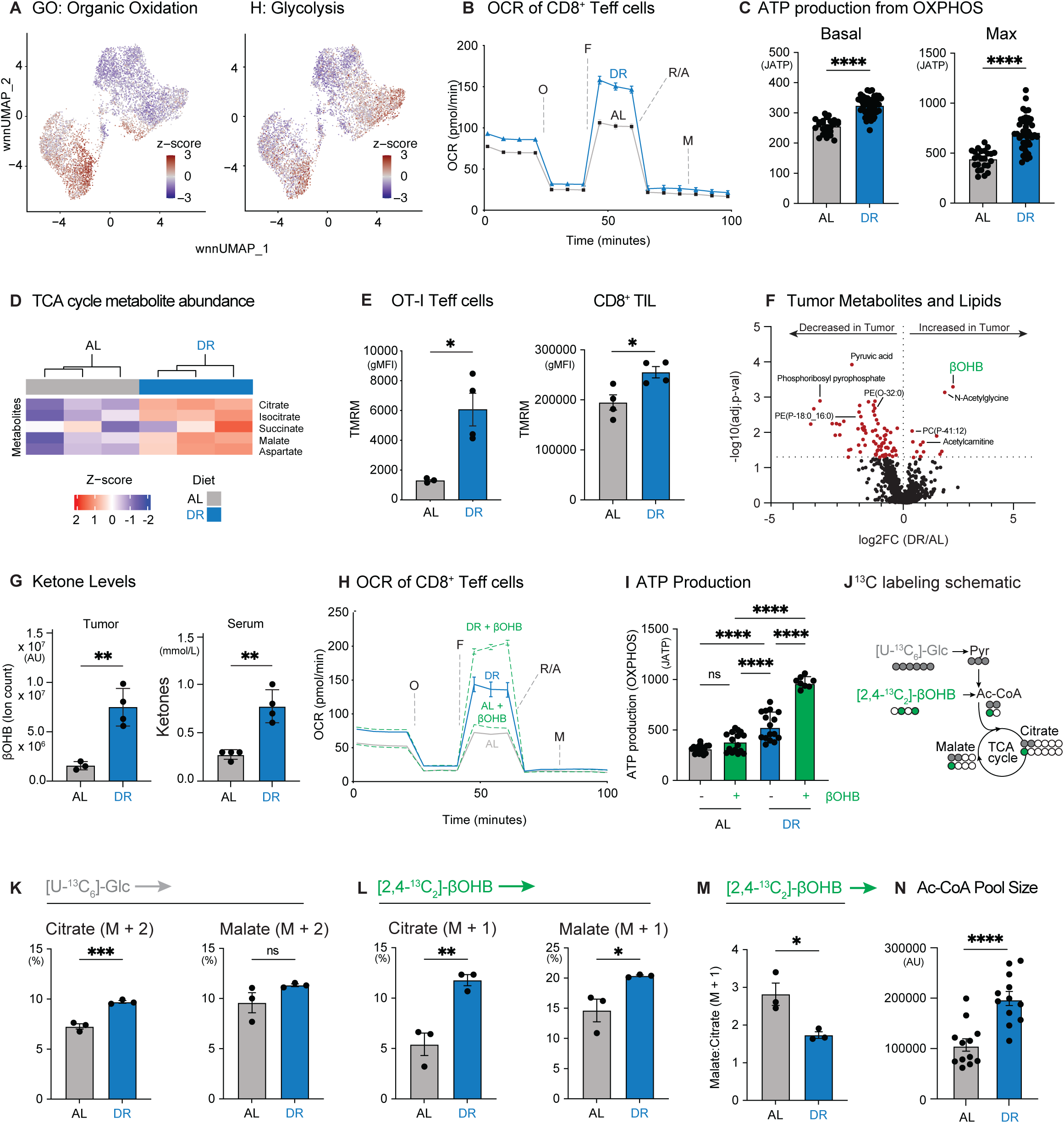
Dietary restriction enhances CD8⁺ T cell metabolic fitness via enhanced ketone body metabolism. A) Overlay of wnnUMAP for 7,005 activated (CD44^+^) CD8⁺ TIL from B16 melanoma tumors (both AL and DR) and MSigDB gene expression signatures for organic oxidation and glycolysis pathways. B-C) Bioenergetic profile of AL- and DR-conditioned CD8⁺ T cells. (B) Oxygen consumption rate (OCR) plot for antigen-specific CD8⁺ T cells isolated from *Lm*OVA-infected AL- or DR-fed mice (7 days post infection (dpi)). Data represent the mean ± SD (n = 24-46, technical replicates). Oligomycin (Oligo), FCCP, rotenone and antimycin A (Rot/AA), and monensin (Mon) were added to cells where indicated. C) Basal and maximal ATP production rates from OXPHOS for AL- or DR-conditioned CD8⁺ T cells from (B). Data represent the mean ± SD (n = 24-46, technical replicates). D) The relative abundance (Z-score) of TCA cycle-derived metabolites in antigen-specific CD8^+^ T cells isolated from *Lm*OVA-infected AL- or DR-fed mice 7 dpi (n=3 mice/group). E) Mitochondrial membrane potential of AL- and DR-conditioned CD8⁺ T cells. *Left,* TMRM staining (gMFI) of CD8^+^ OT-I TIL isolated from B16 tumors from AL- or DR-fed mice (14 days PTI). *Right*, TMRM staining of CD8^+^ OT-I T cells isolated from the spleen of *Lm*OVA-infected AL- or DR-fed mice (7 dpi). Data represent the mean ± SEM (n = 4 mice/group). F) Volcano plot showing the log_2_ fold change in metabolite and lipid abundance in B16 tumors isolated from AL versus DR mice (n = 3-4 mice/group). Select metabolites enriched in AL and DR tumors are annotated. G) Ketone body levels in the serum and tumors of AL-versus DR-fed mice after 21 days on diet (14 days of tumor growth). *Left,* βOHB abundance in B16 tumors from AL-versus DR-fed mice as quantified by mass spectrometry. *Right*, Ketone body abundance in serum as quantified by enzyme assay. Data represent the mean ± SEM (n=3-4 mice/group). H-I) Bioenergetic profile of AL- and DR-conditioned CD8⁺ T cells exposed to βOHB. (H) OCR plot for antigen-specific CD8⁺ T cells isolated from *Lm*OVA-infected AL-or DR-fed mice (7 dpi). T cells were cultured with or without 1.5 mM βOHB 60 minutes prior to the start of the assay. I) Maximum ATP production rates from OXPHOS for CD8⁺ T cells from (H). Data represent the mean ± SD (n = 8-16, technical replicates). J) Schematic depicting ^13^C labeling patterns in acetyl-CoA (Ac-CoA) and TCA cycle intermediates from U-[^13^C₆]-glucose and [2,4-^13^C_2_]-βOHB. K-M) ^13^C labeling of TCA cycle intermediates in AL-versus DR-conditioned CD8⁺ T cells. CD8⁺ T cells isolated from *Lm*OVA-infected AL- or DR-fed mice (7 dpi) were cultured for 2h *ex vivo* in VIM medium containing 5 mM U-[^13^C₆]-glucose and 1.5 mM [2,4-^13^C_2_]-βOHB. Shown is the percent incorporation of (K) U-[^13^C₆]-glucose-derived carbon (M+2) and (L) [2,4-^13^C₂]-βOHB-derived carbon (M+1) into citrate and malate. M) Ratio of [2,4-^13^C_2_]-βOHB-labeled malate (M+1) to citrate (M+1) for T cells from panel (L). Data represent the mean ± SEM (n = 3 mice/group). N) Bar graph of Ac-CoA abundance in CD8⁺ T cells isolated from *Lm*OVA-infected AL- or DR-fed mice (7 dpi). Data represent the mean ± SEM (n = 12 mice/group). *P<0.05, **P<0.01, ***P<0.001, ****P<0.0001.

To evaluate how DR impacts CD8^+^ Teff cell metabolism, we used a model of *Listeria monocytogenes* (*Lm*) infection that elicits robust CD8^+^ Teff cell responses in vivo (*18*, *43*, *44*). Thy1.1^+^ OT-I T cell receptor (TCR) transgenic CD8^+^ T cells were transferred into AL or DR-fed mice, followed by infection with attenuated *Listeria* expressing ovalbumin (*Lm*OVA), and splenocytes extracted 7 days post infection (dpi) for functional analysis (**Fig. S7a**). While OT-I CD8^+^ T cells expanded less in DR-fed mice (**Fig. S7b-d**), they displayed increased effector function, characterized by both an increase in the percentage of IFN-γ-producing T cells (**Fig. S7e-f**) and higher IFN-γ protein production on a per-cell basis (**Fig. S7g**). OT-I T cells from DR-treated animals also displayed a ∼2-fold increase in Granzyme B production (**Fig. S7h**). Metabolically, OT-I Teff cells from DR mice displayed increased rates of oxygen consumption (**Fig. 3b**) and extracellular acidification (**Fig S8a**), corresponding to increased rates of ATP production from both OXPHOS and glycolysis compared to CD8^+^ T cells from AL-fed animals (**Fig. 3c, S8b-c**). Consistent with increased mitochondrial metabolism, steady state levels of TCA cycle-derived metabolites were elevated in DR-conditioned OT-I Teff cells (**Fig. 3d, S8d**). In addition, both OT-I Teff cells responding to *Listeria* infection and CD8^+^ TILs isolated from B16 tumors displayed increased mitochondrial membrane potential under DR conditions compared to CD8^+^ T cells from AL-fed mice (**Fig. 3e**). Together, these data indicate that DR boosts CD8^+^ Teff cell bioenergetics, promoting increased mitochondrial metabolism and OXPHOS.

Dietary modifications that alter systemic metabolism in mice can impact nutrient availability in the TME (*21*, *27*, *45*). Thus, we hypothesized that DR impacts CD8^+^ T cell metabolism in part by altering the availability of specific nutrients in vivo. Indirect calorimetry revealed that mice conditioned to DR displayed a significant decrease in their respiratory exchange ratio (RER) during the fasted−but not the fed−state (**Fig. S8e**), indicating increased lipid oxidation in DR-fed mice. We next used mass spectrometry to profile diet-induced alterations in lipid and metabolite abundance in B16 tumors. We observed a general decrease in fatty acid and lipid abundance in DR tumors compared to AL-fed controls (**Fig. 3f, S8f, Tables S5-6**). This change in tumor lipid abundance is consistent with lipid mobilization from adipose tissue and a decrease in respiratory exchange rate (RER) favoring oxidation triggered by DR (*27*, *33*) (**Fig. S8e**). In contrast, only a small number of metabolites were enriched in tumors from DR-treated mice, including increased abundance of the ketone β-hydroxybutyrate (βOHB) (**Fig. 3f**). Elevated βOHB levels were observed in both tumors and serum from DR mice compared to AL-fed mice (**Fig. 3g**).

The increase in circulating and intra-tumoral βOHB levels under DR was notable given recent evidence linking ketone body (KB) metabolism to enhanced T cell effector function (*19*, *46*). Using Seahorse bioenergetic analysis and ^13^C-labeled metabolic tracers, we found a striking difference in βOHB utilization by CD8^+^ Teff cells based on diet. First, augmenting DR-conditioned OT-I Teff cells with βOHB further increased their maximal oxygen consumption rate (**Fig. 3h**). This change corresponds to a ∼4-fold increase in maximal oxidative ATP production capacity compared to Teff cells from AL-fed animals (**Fig. 3h-i, S8h-g**). Next, we cultured OT-I Teff cells from *Lm*OVA-infected mice ex vivo in physiologic medium (VIM) (*18*) containing fully-labeled [^13^C_6_]-glucose and partially labelled [2,4-^13^C_2_]-βOHB. In this strategy, breakdown of [^13^C_6_]-glucose generates M+2 labeled TCA cycle intermediates whereas [2,4-^13^C_2_]-βOHB generates M+1 labeled intermediates (**Fig. 3j**), which thereby allowed us to directly compare the contribution of each fuel type to TCA cycle metabolism (*19*). We observed a small but significant increase in [^13^C_6_]-glucose-derived citrate (M+2) in DR-conditioned CD8^+^ Teff cells (**Fig. 3k, Table S7**); however, citrate labeling from [2,4-^13^C_2_]-βOHB (M+1) doubled in CD8^+^ Teff cells from DR compared to AL-fed mice (**Fig. 3l**). DR-conditioned OT-I Teff cells also displayed a lower ratio of [2,4-^13^C_2_]-βOHB-labeled malate to citrate (**Fig. 3m**), indicative of increased export of mitochondrial citrate to the cytosol (*47*). Consistent with these observations, steady state acetyl-CoA levels were 2-fold higher in CD8^+^ Teff cells from DR mice relative to T cells from AL-fed controls (**Fig. 3n**). Thus, DR increases both systemic βOHB availability and its utilization by CD8^+^ T cells.

### Ketone bodies fuel anti-tumor immunity under dietary restriction

One of the defining features of CD8^+^ TILs from DR tumors is the expansion of proliferating TOX^+^ Tex cells with effector-like properties (Tex^Eff^ cells, **Fig. 2c-d**). To better understand the metabolic properties of these cells, we analyzed transcriptional profiles of CD8^+^ TIL subsets from human tumors. Similar to CD8^+^ TILs from mouse tumors (**Fig. 2**), we identified CD8^+^ effector T (Teff) cells and two exhausted (Tex) cell clusters in TILs from human tumors (**Fig. 4a**). The dysfunctional or exhausted-like (Tex) cell populations were distinguished from each other by the expression of the proliferation marker *MKI67* (Tex^Eff^ versus Tex^Term^, **Fig. S8i**). Tex^Eff^ cells from human tumors most closely associated with Tex^Eff^ cluster of cells from mouse TILs (**Fig. 2a**). Notably, both human Teff and Tex^Eff^ cells displayed significant increases in transcript levels for βOHB dehydrogenase (*BDH1*), which encodes the rate-limiting enzyme required for βOHB breakdown (**Fig. 4b**). Interestingly, proliferating Tex^Eff^ cells (human TIL) displayed the highest ketone body metabolism gene signature of all human TIL subsets (**Fig. 4c**), driven by increased expression of *BDH1* and 3-oxoacid CoA-transferase 1 (*OXCT1*), which converts βOHB to acetoacetate (AcAc) (**Fig. 4b**).

**Fig. 4.**
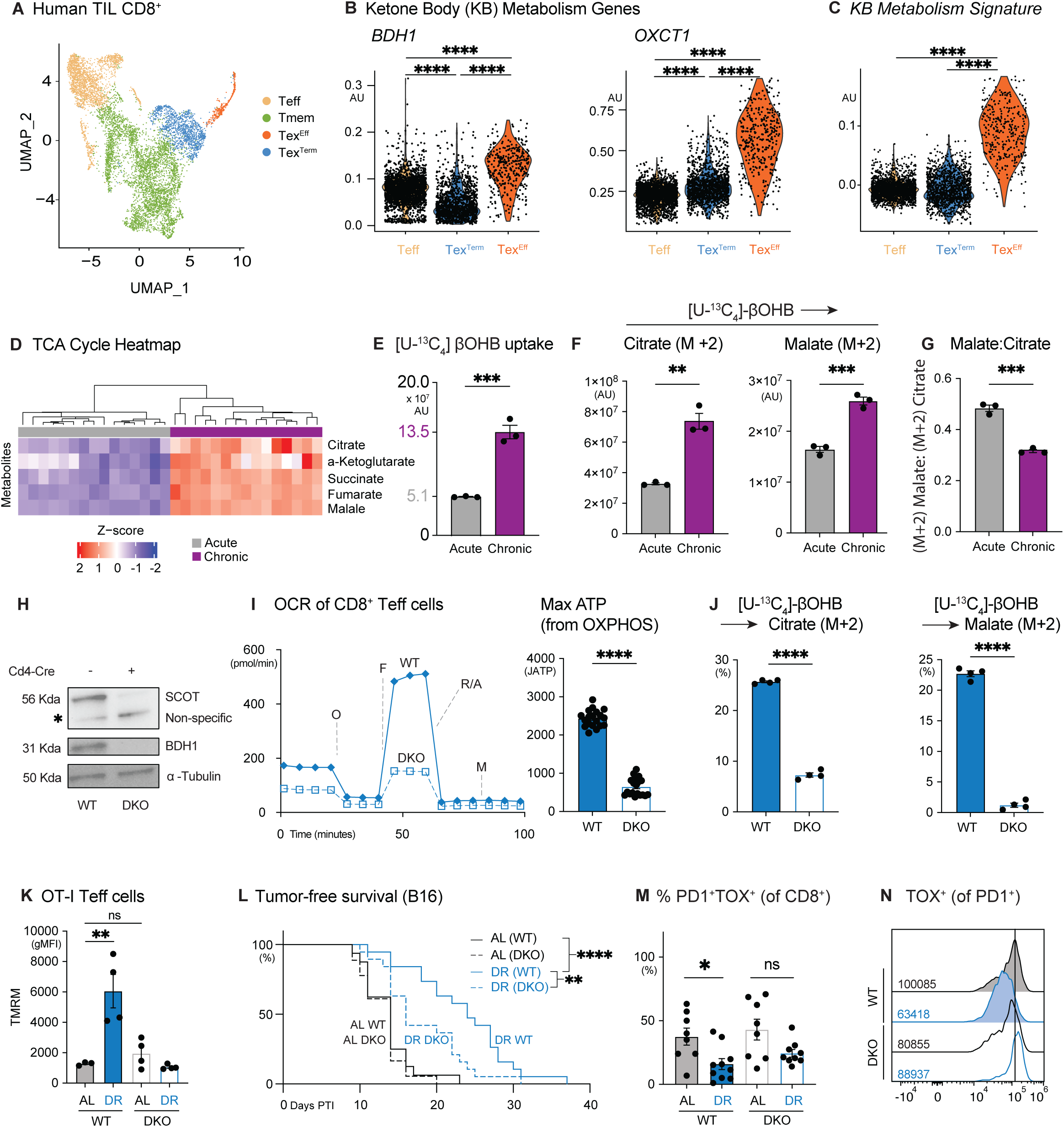
Ketone bodies fuel anti-tumor immunity under dietary restriction. A) Uniform Manifold Approximation and Projection (UMAP) of 8,552 cells human CD8⁺ tumor-infiltrating lymphocytes (TILs) from GSE98638. Shown are unique clusters for effector (Teff), memory (Tmem), terminally exhausted (Tex^Term^), and proliferating exhausted (Tex^Eff^) CD8⁺ T cell populations. B-C) Violin plots of gene expression across human CD8⁺ TIL subsets. B) Expression of *BDH1*, and *OXCT1* genes. C) Expression of ketolysis signature genes (Ketolysis gene set: *ACAT1*, *ACAT2*, *BDH1*, *BDH2*, *HMGCL*, *HMGCS1*, *HMGCS2*, *OXCT1*, *OXCT2*). Statistical significance was assessed by one-way ANOVA with Dunnett’s multiple comparisons test and a 5% significance level. D) Heatmap showing the relative abundance (Z-score) of TCA cycle-derived metabolites from CD8⁺ T cells exposed to acute versus chronic stimulation with anti-CD3 and -CD28 antibodies *in vitro* (n = 3 mice/group). E) Bar graph depicting total abundance of ^13^C-labeled βOHB in acute versus chronic stimulated CD8⁺ T cells as in (D). Data represent the mean ± SEM (n = 3 mice/group). F-G) ^13^C labeling of TCA cycle intermediates in CD8⁺ T cells exposed to chronic antigen stimulation. CD8⁺ T cells exposed to acute versus chronic stimulation as in (D) were cultured in VIM medium containing 0.85 mM [^13^C_4_]-βOHB for 2h. F) Total abundance of [^13^C4]-βOHB-derived citrate and malate. G) Ratio of [^13^C_4_]-βOHB-labeled malate (M+2) to citrate (M+2) for T cells from (F). Data represent the mean ± SEM (n = 3 mice/ group). H) Immunoblot of SCOT and BDH1 protein expression in CD8⁺ T cells from wild type (WT, *Cd4-Cre*^-^; *Bdh1*^fl/fl^*Oxct1*^fl/fl^) or *Bdh1/Oxct1* double knockout (DKO, *Cd4-Cre*^+^; *Bdh1*^fl/fl^*Oxct1*^fl/fl^). Protein levels of α-tubulin are shown as a loading control. I) Bioenergetic profile of control (WT) and ketolysis-deficient (DKO) CD8⁺ T cells under DR conditions. *Left,* OCR plot of WT and DKO OT-I T cells isolated from DR-fed *Lm*OVA-infected mice (7 dpi). *Right*, bar graph showing maximal ATP production rates from OXPHOS for WT versus DKO CD8⁺ T cells. Data represent the mean ± SM (n=19-20). J) ^13^C labeling of TCA cycle intermediates in control (WT) and ketolysis-deficient (DKO) CD8⁺ T cells under DR conditions. Bar graph showing U-[^13^C4]-βOHB labeling in citrate M+2 (*left*) and malate M+2(*right*) in WT versus DKO CD8⁺ T cells isolated from DR-fed *Lm*OVA-infected mice (7 dpi). Data represent the mean ± SEM (n = 4 mice/group). K) Mitochondrial membrane potential of control (WT) versus ketolysis-deficient (DKO) CD8⁺ T cells under DR conditions. Bar plot showing TMRM staining of CD8⁺ T cells isolated from *Lm*OVA-infected AL- or DR-fed mice (7 dpi). Data represent the mean ± SEM (n = 4 mice/group). L) Kaplan-Meier plot comparing tumor onset (tumor volume ≥250 mm³) in B16 tumor-bearing WT versus DKO mice fed an AL or DR diet as in Figure 1D. Statistical significance was assessed by log-rank test with Bonferroni correction (n=16-19 mice/group). M-N) Enhanced exhaustion of DKO CD8⁺ T cells under DR-fed conditions. (M) Bar plot showing the percentage of PD1⁺TOX⁺ CD8⁺ T cells isolated from B16 tumors from WT versus DKO mice under AL- or DR-fed conditions. N) Representative histograms of TOX expression in CD8⁺ TIL isolated from B16 tumors from WT and DKO mice fed under AL- or DR-conditions. *Inset,* gMFI values for TOX expression averaged across all biological replicates. Data represent the mean ± SEM (n = 8-10). *P<0.05, **P<0.01, ***P<0.001, ****P<0.0001.

In light of these findings, we assessed the metabolism of CD8^+^ T cells exposed to chronic antigenic stimulation that limits their self-renewal capacity and promotes terminal differentiation (**Fig. S9a**) (*17*). Chronic stimulation of in vitro-activated CD8^+^ T cells with anti-CD3 and -CD28 antibodies promotes several features of T cell exhaustion including increased expression of TOX and inhibitory receptors (i.e., PD1, TIM3) (**Fig. S9b**), as well as reduced polyfunctionality (**Fig. S9c**) compared to activated CD8^+^ T cells maintained in IL-2 (acute stimulation). Surprisingly, chronically stimulated CD8^+^ T cells displayed increased abundance of TCA cycle intermediates compared to acutely stimulated controls (**Fig. 4d**). This increase in TCA cycle intermediates was similar to DR-conditioned CD8^+^ T cells (**Fig. 3d**), leading us to hypothesize that chronic antigen exposure may promote increased KB utilization. Using ^13^C-βOHB as a metabolic tracer, we found that chronically stimulated CD8^+^ T cells displayed increased βOHB uptake (**Fig. 4e, Table S8**) as well as higher βOHB oxidation in the TCA cycle as determined by increased levels of ^13^C-βOHB-derived citrate and malate compared to controls (**Fig. 4f, Table S8**). Chronically stimulated CD8^+^ T cells also displayed a reduced malate-to-citrate ratio (**Fig. 4g, Table S8**). This suggested a net export of βOHB-derived citrate to the cytosol, similar to our observations for DR-conditioned T cells (**Fig. 3m**). Collectively, these data suggest that chronic antigen exposure stimulates increased βOHB uptake and metabolism by CD8^+^ T cells.

Next, we assessed the contribution of βOHB metabolism by T cells to the anti-tumor effects of DR. To test this, we generated mice with T cell-specific deletion of the enzymes required to process KBs, βOHB dehydrogenase 1 (BDH1) and succinyl-CoA:3-ketoacid CoA transferase (SCOT; encoded by *Oxct1*), by crossing *Cd4*^Cre^ transgenic mice to mice with floxed alleles targeting the *Bdh1* and *Oxct1* genes (Bdh1^fl/fl^Oxct1^fl/fl^*Cd4-Cre*, **Fig. S10a**). BDH1 mediates the conversion of βOHB to acetoacetate (AcAc), while SCOT converts AcAc to acetoacetyl-CoA, ultimately leading to the production of acetyl-CoA destined for the TCA cycle (**Fig. S10b**). Double knockout (DKO) T cells (from *Bdh1*^fl/fl^*Oxct1*^fl/fl^*Cd4-Cre* mice) lacking both BDH1 and SCOT were confirmed by immunoblot (**Fig. 4h**). Seahorse analysis of control (WT, from Cre-negative *Bdh1*^fl/fl^*Oxct1*^fl/fl^ mice) or DKO OT-I CD8^+^ T cells isolated from *Lm*OVA-infected mice demonstrated that loss of BDH1 and SCOT reversed the boost in OXPHOS induced by DR feeding (**Fig. 4i, Fig. S10c-d**). Consistent with these observations, DKO CD8^+^ OT-I cells isolated from DR-fed mice displayed a significant reduction in ^13^C-βOHB labeling of TCA cycle intermediates (i.e., citrate, malate) compared to controls (**Fig. 4j**). TMRM staining also revealed that the DR-induced increase in mitochondrial membrane potential was blunted in DKO CD8^+^ T cells (**Fig. 4k**). Together, these data indicate that T cell-intrinsic ketolysis is responsible for maintaining T cell mitochondrial function and bioenergetic capacity under DR.

Finally, we assessed the contribution of T cell-intrinsic ketolysis to the anti-tumor effects of DR by challenging control (WT) or DKO mice with syngeneic tumors. We observed no difference in B16 melanoma tumor growth between DKO and control mice under AL conditions; however, tumor growth was accelerated in DKO mice specifically under DR conditions (**Fig. 4l**). Analysis of CD8^+^ TIL from these mice revealed increases in the accumulation of PD1^+^TOX^+^ Tex cells in the tumors of DR-fed DKO mice compared to control animals (**Fig. 4m**). Furthermore, these DKO CD8^+^ TIL maintained high TOX levels even under DR feeding conditions (**Fig. 4n**). Together, these data link T cell-intrinsic ketolysis to the anti-tumor effects of DR, with ketolysis deficient T cells displaying increased features of exhaustion (i.e., TOX expression) in the TME.

## Discussion

The tumor suppressive effects of reduced calorie intake have long been presumed to act directly on cancer cells. Here, we demonstrate that dietary restriction (DR) works more broadly to limit tumor growth by stimulating CD8^+^ T cell anti-tumor immunity. We show that DR alters CD8^+^ T cell fate in the TME, promoting the expansion of tumor-controlling effector (Teff-like and Tex^Eff^) cells while limiting terminal T cell exhaustion. Moreover, DR enhances CD8^+^ T cell anti-tumor immunity by increasing circulating KB levels, which in turn enhance TCA cycle metabolism and mitochondrial bioenergetics of CD8^+^ T cells. T cells that cannot metabolize KBs display metabolic deficits, undergo premature exhaustion, and fail to control tumor growth under DR conditions, identifying T cell-intrinsic ketolysis as a central mechanism for the anti-tumor effects of DR. These findings, along with our previous research identifying ketolysis as a regulator of CD8^+^ T cell cytolytic function (*19*, *46*), suggest that DR regulates a *nutrient-sensitive checkpoint* within the TME−mediated by ketones−that promotes the expansion of tumor-controlling effector (Teff-like and Tex^Eff^) T cells over terminal exhaustion, thereby improving tumor control. Overall, our study highlights how altering systemic nutrient availability through diet can influence CD8^+^ T cell fate within tumors to limit cancer progression.

βOHB is preferentially oxidized over glucose for ATP production in CD8^+^ T cells and boosts T cell effector responses (*21*, *48*). Our data indicate that βOHB uptake and oxidation is a metabolic feature of T cells exposed to chronic stimulation. Increased *BDH1* expression in CD8^+^ T cells, which we observed in TILs from human tumors, facilitates greater βOHB oxidation in TIL as they infiltrate solid tumors. Many tumors are nutrient-poor (*19*). By increasing KB availability approximately fourfold compared to *ad libitum*-fed conditions, DR boosts the available βOHB supply in the TME, fueling the TCA cycle to meet the oxidative demands of CD8^+^ TIL. Consistent with this, CD8^+^ T cells adapted to DR were found to display increased mitochondrial membrane potential and oxidative ATP production. This metabolic advantage promotes the expansion of Tex^Eff^ cells, which retain proliferative capacity and effector function, over Tex^Term^ cells with high inhibitory receptor expression and diminished functionality. These metabolic advantages are lost in ketolysis-deficient (DKO) T cells, contributing to their premature exhaustion. Thus, DR aligns systemic nutrient supply with the metabolic needs of T cells at the tissue level.

A critical metabolic fate of βOHB oxidation in CD8^+^ T cells is acetyl-CoA, the levels of which double under DR-fed conditions. We have shown that βOHB is the major substrate for acetyl-CoA production in CD8^+^ T cells under physiologic conditions (*49*, *50*). The role of acetyl-CoA extends beyond biosynthetic growth (i.e., *de novo* lipogenesis) and energy production: it is the rate-limiting substrate for histone acetylation reactions that regulate T cell differentiation and effector function (*20*). Our results here argue that nutritional regulation of KB metabolism is a critical determinant of CD8⁺ T cell fate in the TME, shifting differentiation between terminally exhausted (Tex^Term^) and effector-like (Tex^Eff^) states. Consistent with this, pantothenate/Coenzyme A (CoA) increases CD8⁺ T cell differentiation towards effector lineages to enhance tumor control (*51–53*). We speculate that, under DR, βOHB-dependent changes in acetyl-CoA levels drive changes in T cell epigenetic programming favoring effector responses over exhaustion. Thus, dietary interventions or therapeutics that boost acetyl-CoA production in T cells may improve anti-tumor immunity similar to DR.

The mammalian immune system evolved under cyclical periods of feast and famine. Immune challenges such as infection further impact systemic nutrient availability by disrupting feeding behaviors and remodeling host metabolism (*54*, *55*). Mobilizing stored energy from adipose tissue into KB production provides the host with a versatile fuel to maintain acetyl-CoA production under conditions of metabolic stress (*56*). In this vein, we speculate that T cells evolved the use of ketolysis to buffer against metabolic perturbations that negatively impact T cell bioenergetics and function. Our findings reveal the potential of exploiting this system through nutritional interventions that enhance anti-tumor immunity. PD1 blockade amplifies the anti-tumor effects of DR by promoting the expansion of effector T cells (Teff and Tex^Eff^ cells) to limit tumor growth, highlighting the potential of enhancing the efficacy of immunotherapies through nutritional intervention. Employing DR in clinical settings may face challenges regarding feasibility due to patient health and compliance. However, combining ICIs with pharmacological agents that reduce appetite and food intake, such as GLP-1 agonists like semaglutide (i.e., Ozempic), may mimic some metabolic effects of DR without necessitating strict dietary regimens (*57*). Our findings also have potential implications for adoptive T cell therapies, including chimeric antigen receptor (CAR) T-cell therapy. The metabolic state of T cells prior to infusion is critical for their persistence, functionality, and anti-tumor efficacy in patients (*45*). Exposing CAR T cells to a DR-like environment during expansion or enhancing their capacity to oxidize KBs may improve their metabolic fitness and resistance to terminal exhaustion *in vivo*, thereby increasing their therapeutic efficacy.

One caveat of this work is that the chow used is relatively low-fat (15% of calories from fat) compared to the standard western diet. High-fed feeding antagonizes anti-tumor T cell responses (*58*); thus, it is crucial to dissect the impact of macronutrient content, particularly carbohydrate and fats (*19*), on the anti-tumor effects of DR. Understanding how dietary interventions such as DR impact T cell metabolism and differentiation fate in the TME may lead to evidence-based nutritional guidelines that complement and enhance cancer immunotherapy efforts.

## Acknowledgements

We acknowledge Drs. Evan Lien, Julian Lum, Tim Triche, Bart Williams, and members of the Jones and Krawczyk laboratories for scientific discussions contributing to this manuscript. We thank Noah Lubben for technical assistance and Jeanie Wedberg, Margene Brewer, and Michelle Minard for administrative assistance. We thank members of the VAI Core Facilities (Mass Spectrometry, Bioinformatics and Biostatistics, Flow Cytometry, and Vivarium) for technical assistance.

## Funding

VAI Metabolism & Nutrition (MeNu) Program Pathway-to-Independence Award (to JL)

Canadian Institutes of Health Research (CIHR) Fellowship (MFE-181903, to JL)

National Institutes of Health (NIH)/National Cancer Institute (NCI)

T32 training grant (T32CA251066-01A1, to MJW)

Damon Runyon Foundation Postdoctoral Fellowship (2495-23, to MJW)

NIH/National Institute of Allergy and Infectious Diseases (NIAID) R21 award (R21AI153997, to CMK)

Cancer Research Institute fellowship (to SM)

Salk Pioneer Fund Postdoctoral Scholar Award (to SM).

NIH/NIAID R01 award (R01AI066232, to SMK)

NIH/NIAID R21 award (R21AI151986, to SMK).

Paul G. Allen Frontiers Group Distinguished Investigator Program (to RGJ)

NIH/NIAID R01 award (R01AI165722, to RGJ)

## Author Contributions

Conceptualization: BMO, LMD, RGJ

Methodology: BMO, LMD, RDS, MTSH, MPV, BKJ, HJJ, CMK, RGJ

Investigation: BMO, LMD, JL, NB, MSD, NB, SM, MJW, AEE, CNI

Visualization: BMO, SM, BKJ, KSW

Funding acquisition: SK, RGJ

Project administration: RGJ

Supervision: RDS, HS, PAC, SK, CMK, RGJ

Writing – original draft: BMO, RGJ

Writing – review & editing: BMO, JL, RGJ

## Data Availability Statement

All unique and stable reagents generated in this study are available from the Lead Contact upon completion of a Materials Transfer Agreement. CITE-sequencing data of mouse tumor-infiltrating lymphocytes have been deposited in the National Center for Biotechnology Information Gene Expression Omnibus (NCBI GEO) under accession number <u>GSE267070</u>. Human single-cell RNA-sequencing data of tumor-infiltrating lymphocytes used in this research are available at NCBI GEO under accession number <u>GSE146771</u>. All other data supporting the findings of this study are available within the article and its online supplementary material. Bioenergetics data were analyzed using protocols developed by Mookerjee and Brand, available for download at https://russelljoneslab.vai.org. R code used for data processing is available at our GitHub repository (https://github.com/rgjcanada/DR_CITESEQ_2025.git). Processed data files (RDS files) curated from the data processing can be found at https://doi.org/10.5281/zenodo.13920173.

## Supplementary Materials

Materials and Methods

Figs. S1 to S10

Tables S1 to S8

