## Supplemental Text/Methods for "Dietary Restriction Enhances CD8⁺ T Cell Ketolysis to Limit Exhaustion and Boost Anti-Tumor Immunity"

**The PDF file includes:**

Materials and Methods
Figs. S1 to S10
References

#### Materials and Methods

##### Mice

This study used the following mouse strains: C57BL/6J (RRID: IMSR_JAX:000664); B6.PL-*Thy1*$^{a}$/CyJ (Thy1.1; RRID: IMSR_JAX:000406), B6.SJL-Ptprca Pepcb/BoyJ (CD45.1+), Tg(TcraTcrb)1100Mjb (OT-I; RRID: IMSR_JAX:003831). The mice were ordered from The Jackson Laboratory. The *Bdh1*$^{fl/fl}$/*Oxct1*$^{fl/fl}$*Cd4-Cre* line was generated by crossing *Bdh1*$^{fl/fl}$*Cd4-Cre* mice (*1*) and *Oxct1*-floxed mice (provided by Peter Crawford (*2*, *3*)). *Bdh1*$^{fl/fl}$*Oxct1*$^{fl/fl}$*Cd4-Cre* OT-I mice were generated by crossing *Bdh1*$^{fl/fl}$*Oxct1*$^{fl/fl}$*Cd4-Cre* mice with the Tg(TcraTcrb)1100Mjb mouse line. All mice were bred and housed under specific pathogen-free conditions at VAI, following approved protocols. Genotyping was conducted using DNA extracted from tail or ear biopsies, with primer sets listed in Key Resources Table. The study included both male and female mice aged 8 to 14 weeks.

##### Cell lines

B16-F10 murine melanoma cells expressing OVA (B16-OVA; (*4*)) and EO771 breast cancer cells (CRL-3461) were cultured in Dulbecco's Modified Eagle's Medium (DMEM) from Wisent Inc., supplemented with 10% heat-inactivated fetal bovine serum (FBS), 1% penicillin-streptomycin (Gibco), and 6 mM L-glutamine. All cell cultures were maintained in a humidified incubator at 37°C with 5% $CO_2$.

##### Tumor models

Male and female C57BL/6J, *Bdh1*$^{fl/fl}$*Oxct1*$^{fl/fl}$ (wild-type; WT), or *Bdh1*$^{fl/fl}$*Oxct1*$^{fl/fl}$*Cd4-Cre* (double knockout; DKO) mice were maintained a 5010 diet. Mice fed *ad libitum* (AL) were allowed free access to food at all times and never ran out of food. The mice on the DR regimen received 50% of their average daily (normal) intake of food. Mice on DR were given pre-weighed pellets between 9 – 11 am for every day of the experiment. Following a week on their respective diet, mice were subcutaneously injected with 2.5 x $10^5$ (EO771) or 5 x $10^5$ (B16-OVA) cells into the right abdominal flank. Tumor volume was measured every 2-3 days with a caliper once tumors became palpable. Tumor initiation was scored as a tumor volume ≥250 $mm^3$. Mice were euthanized when tumor volume reached 1500 $mm^3$. For experiments involving anti-PD1 treatment, mice received 200 µg of IgG (BP0091; RRID: AB_1107773) or anti-PD1 (BP0033-2; RRID:AB_1107747) antibodies intraperitoneally every 3 days, for a total of 5 injections (1 mg of antibody total). Treatment was initiated 7 days after tumor cell injection.

**Tumor infiltrating lymphocyte (TIL) isolation**

TIL were isolated from palpable tumors 12-14 days post-tumor cell injection. Tumors were mechanically homogenized in a 6-well plate and then passed through a 100 µm cell strainer, followed by a 40 µm cell strainer. The single-cell suspension was incubated with red blood cell (RBC) lysis buffer for 1 minute at room temperature, after which three volumes of T cell media (TCM) were added to halt the lysis reaction. Cells were collected by centrifugation at 500 RCF for 5 minutes at 4°C, and then resuspended in 0.5-1 mL of TCM and processed for flow cytometry.

**Mouse T Cell Isolation and Culture**

$CD8^+$ T cells were purified from mouse spleens through negative selection using magnetic bead-based isolation kits (StemCell Technologies). T cells were cultured in T cell medium (TCM): Iscove's Modified Dulbecco's Medium (IMDM; Wisent Inc.) containing 10% Nu-Serum IV culture supplement (Corning), 50 U/mL penicillin, 50 µg/mL streptomycin (Gibco), 50 µM 2-mercaptoethanol (Gibco), and 2 mM L-glutamine (Gibco; final concentration 6 mM). T cells ($1 \times 10^6$ cells/mL) were activated in vitro by stimulation with plate-bound anti-CD3ε (clone 145-2C11; 2 µg/mL) and anti-CD28 (clone 37.51; 1 µg/mL) antibodies for 48-72 hours. After activation, cells were cultured in IMDM with 50 U/mL recombinant murine IL-2 (Peprotech) and re-seeded at $4 \times 10^5$ cells/mL in fresh medium supplemented with IL-2 every two days.

For induction of in-vitro chronically stimulated cells, $CD8^+$ T cells were activated in vitro for 48 hours as described above, followed by culture on fresh plate-bound anti-CD3ε and anti-CD28 antibodies for an additional 48 hours as previously described (*5*). Acutely activated cells were maintained in IL-2 (50 U/ml) without restimulation.

**Adoptive Transfer and *L. monocytogenes* (*Lm*OVA) Infection**

Mice were intravenously injected with a sublethal dose of recombinant *Listeria monocytogenes* expressing OVA (*Lm*OVA, $2 \times 10^6$ CFU), following a protocol previously established (*1*). For adoptive transfer studies involving analysis on days 6-7 post-infection (dpi), $5 \times 10^3$ OT-I $CD8^+$ T cells ($Thy1.1^+$ or $CD45.2^+$) were transferred intravenously into C57BL/6J recipients ($Thy1.2^+CD45.2^+$ or $CD45.1^+$) on day -1, and *Lm*OVA infection was induced 24 hours later (day 0). Splenocytes were collected 7 dpi to evaluate OVA-specific $CD8^+$ T cells through Thy1.1 or CD45.2 staining. Cytokine production was measured using intracellular cytokine staining (ICS) following re-stimulation with OVA peptide ($OVA_{257–264}$) for 4 hours (GolgiStop added at 1ug/ml after 2h of restimulation) at 37°C.

For metabolic analysis of *Lm*OVA-specific $Thy1.1^+$ OT-I T cells *ex vivo*, $Thy1.2^+$ C57BL/6 mice received 5 x $10^4$ $Thy1.1^+$ OT-I T cells at day -1, followed by *Lm*OVA infection on day 0. *Lm*OVA-specific $CD8^+$ OT-I T cells were isolated from the spleen of infected mice by positive selection using the EasySep mouse CD90.1 positive selection kit (StemCell Technologies) as previously described (*6*, *7*). For ex vivo $^{13}C$ isotope tracing, 5 × $10^4$ $Thy1.1^+$ OT-I $CD8^+$ T cells were injected into $Thy1.2^+CD45.2^+$ C57BL/6J mice, followed by *Lm*OVA infection the next day. At 7 dpi, activated $Thy1.1^+$ OT-I T cells were isolated from the spleens using magnetic bead purification (*6*, *7*) and cultured in vitro for 2h in VIM medium (*8*) containing 1.5 mM βOHB.

**Ketone body measurements**

To monitor blood ketone levels, blood was collected via the submental vein from restrained mice and ketone test strips (Keto-Mojo) used to quantify blood ketone levels.

**Metabolite and lipid extraction**

For T cell tracing and metabolite profiling studies, metabolites were extracted by mixing with ice cold acetonitrile:methanol:water (4:4:2, v/v) (*9*), sonicating for 5 min, and incubating on wet ice for 1h. Extracts were centrifuged and the soluble fraction was collected and dried under vacuum. Extracts were resuspended 50 μL of water for LC-

MS analysis. For tissue and plasma metabolite and lipid profiling, samples were extracted with chloroform:methanol:water (2:2:1.8 v/v) (*9*, *10*). For plasma, this was accomplished by mixing 40 μL of plasma with 690 μL 1:1 chloroform methanol. For tissue, 690 μL of 1:1 chloroform:methanol was added to 40 mg of tissue and homogenized with a beadmill homogenizer. For both sample types, 310 μL of water was added, incubated on wet ice for one hour and centrifuged at 14000g for 10 minutes to induce phase separation. 495 μL and 100 μL of the upper aqueous and bottom organic layers, respectively, were collected into separate tubes and dried in a rotary vacuum evaporator. The aqueous layer was resuspended 100 μL of water for metabolomics analysis. The organic layer was resuspended 200 μL of 50:50 (isopropanol:acetonitrile, v/v) for lipidomics analysis.

**Metabolomics Analyses**

Metabolomic profiling and stable isotope tracing data was collected using a Vanquish liquid chromatography system coupled to an Orbitrap Exploris 240 (Thermo Fisher Scientific) using an H-ESI (heated electrospray ionization) source in negative mode as previously described (*1*, *9*). 2 μL of each standard and/or sample was injected and run through a 24-minute reversed-phase chromatography Zorbax RRHD extend-C18 Column (1.8 μm, 2.1mm × 150mm, 759700-902, Agilent) combined with a Zorbax extend-C18 guard column (1.8 μm, 2.1 mm × 5 mm, 821725-907, Agilent). Mobile phase A consisted of LC/MS grade water (W6, Fisher) with 3% LC/MS grade methanol (A456, Fisher), mobile phase B was LC/MS grade methanol and both mobile phases contained 10 mM tributylamine (90780, Sigma), 15 mM LC/MS grade acetic acid

(A11350, Fisher), and 0.01% medronic acid (v/v, 5191-4506, Agilent). For the wash gradient, mobile phase A was kept the same, and mobile phase B was 99% LC/MS grade acetonitrile (A955, Fisher). Column temperature was kept at 35 °C, flow rate was held at 0.25 mL/min, and the chromatography gradient was as follows: 0-2.5 min held at 0% B, 2.5-7.5 min from 0% B to 20% B, 7.5-13 min from 20% B to 45% B, 13-20 min from 45% B to 99% B, and 20-24 min held at 99% B. A 16 minute wash gradient was run in reverse flow direction between every injection to back-flush the column and to re-equilibrate solvent conditions as follows: 0-3 min held at 100% B and 0.25 mL/min, 3-3.5 min held at 100% B and ramp to 0.8 mL/min. 3.5-7.35 min held at 100% B and 0.8 mL/min, 7.35-7.5 held at 100% B and ramp to 0.6 mL/min, 7.5-8.25 from 100% B to 0% B and ramp to 0.4 mL/min, 8.25-15.5 min held at 0% B and ramp to 0.25 mL/min, and 15.5-16 min held at 0% B and 0.25 mL/min. Mass spectrometer parameters were: source voltage -2500V, sheath gas 60, aux gas 19, sweep gas 1, ion transfer tube temperature 320°C, and vaporizer temperature 250°C. Full scan data were collected using the orbitrap with a scan range of 70-850 m/z at a resolution of 240,000 and RF lens at 35%. Fragmentation was induced in the orbitrap using assisted higher-energy collisional dissociation (HCD) collision energies at 15, 30, and 45%. Orbitrap resolution was 15,000, the isolation window was 2 m/z, and data dependent scans were capped at 5 scans.

For acetyl-CoA measurements, samples were analyzed with a Vanquish liquid chromatography system coupled to an Orbitrap ID-X (Thermo Fisher Scientific) using an H-ESI (heated electrospray ionization) source in positive mode. 2 μL of each standard and/or sample was injected and run through a 5-minute reversed-phase chromatography Cortecs T3 Column (1.6 µm, 2.1 mm x 150 mm, 186008500, Waters) combined with a Cortecs T3 VanGuard Pre-column (1.6 µm, 2.1 mm x 5 mm, 186008508, Waters). Mobile phase A consisted of 100% LC/MS grade water (W6, Fisher), 0.01% ammonium hydroxide (A470, Fisher), 5mM ammonium acetate (73594, Sigma), and mobile phase B consisted of 99% LC/MS grade acetonitrile (A955, Fisher). Column temperature was kept at 30 °C, flow rate was held at 0.3 mL/min, and the chromatography gradient was as follows: 0-0.5 min held at 0% B, 0.5-1 min from 0% B to 10% B, 1-4 min from 10% B to 50% B, 4-4.1 min from 50% B to 99% B, and 4.1-5 min held at 99% B. A 5 minute wash gradient was run between every injection to flush the column and to re-equilibrate solvent conditions as follows: 0-2 min held at 100% B, 2-3 min from 100% B to 0% B, and 3-5 min held at 0% B. Mass spectrometer parameters were: source voltage 3500V, sheath gas 70, aux gas 25, sweep gas 1, ion transfer tube temperature 300°C, and vaporizer temperature 250°C. A targeted single ion scan (tSIM) was performed in the orbitrap to target acetyl-CoA and all of its carbon-13 isotopologues with a center mass of 810.133 m/z and an isolation window 48 m/z. Resolution was set at 60,000, RF lens at 60%, and scan time was set for 2-4 minutes of the chromatographic gradient described above. Data dependent MS2 fragmentation was induced in the orbitrap using assisted higher-energy collisional dissociation (HCD) collision energies at 20, 40, 60, 80, and 100% as well as with collision-induced dissociation (CID) at a collision energy of 35%. For both MS2 fragmentations, orbitrap resolution was 30,000, the isolation window was 1 m/z for HCD and 1.5 m/z for CID, and total cycle time was 0.6 sec.

For βOHB quantification in serum, liver, and spleen, metabolites were extracted with a 40% acetonitrile, 40% methanol, and 20% water solution. An external standard curve for βOHB was generated from 10 µg/mL to 0.01 µg/mL via half-log serial dilutions. Standards were processed identically to tissue and serum samples throughout the workflow. Tissues were extracted at a concentration of 40 mg tissue/mL, and serum at 40 µL serum/mL of extraction solvent. Extracts were dried and reconstituted in water (200 µL for tissue, 1 mL for serum) containing 100 ng/mL of internal standard ([U-$^{13}C_4$]-βOHB). Samples and standards were analyzed on an Orbitrap Exploris 240 (Thermo) in ESI-negative mode using tributylamine ion-paired chromatography. Data were processed using Skyline, and internal standard-normalized peak areas were used to calculate βOHB concentrations, which were reported as mg/g for tissues and mM for serum

**Lipidomics Analyses**

Lipidomics samples were analyzed with a Thermo Vanquish dual liquid chromatography system utilizing two alternating methods, referred to as Chromatography 1 and Chromatography 2, coupled to an Orbitrap ID-X (Thermo Fisher Scientific) using an H-ESI (heated electrospray ionization) source in positive and negative mode respectively. 2 µL of each standard and/or sample was injected, column temperatures were kept at 50 °C, and flow rate was held at 0.4 mL/min. For both chromatography 1 and 2, mobile phase A consisted of 60% LC/MS grade acetonitrile (A955, Fisher Scientific), 40% LC/MS grade water (W6, Fisher Scientific), 0.1% LC/MS grade formic acid (A117, Fisher Scientific), 10mM ammonium formate (70221, Fisher Scientific), and mobile phase B consisted of 90% LC/MS grade isopropanol (A461, Fisher Scientific), 8% LC/MS grade acetonitrile, 2% LC/MS grade water, 0.1% LC/MS grade formic acid, and 10mM ammonium formate. Chromatography 1 used a 30-minute reversed-phase chromatography Accucore C30 column (2.6 µm, 2.1 mm x 150 mm, 27826-152130, Thermo Fisher Scientific) combined with an Accucore C30 guard column (2.6 µm, 2.1mm x 10 mm, 27826-012105, Thermo Fisher Scientific), and the gradient was as follows: 0-1 min held at 25% B, 1-3 min from 25% B to 40% B, 3-19 min from 40% B to 75% B, 19-20.5 min 75% B to 90% B, 20.5-28 min from 90% B to 95% B, 28-28.1 min from 95% B to 100% B, and 28.1-30 min held at 100% B. A 30 minute wash gradient was run between every injection (in parallel with chromatography 2) to flush the column and to re-equilibrate solvent conditions as follows: 0-2 min held at 100% B and 0.3 mL/min, 2-2.1 min from 100% B to 25% B and held at 0.3 mL/min, 2.1-4 min held at 25% B and ramp to 0.4 mL/min, 4-6 held at 25% B and ramp to 0.6 mL/min, 6-17 min held at 25% B and 0.6 mL/min, 17-17.1 min held at 25% B and ramp to 0.4 mL/min, and 17.1-30 min held at 25% B and 0.4 mL/min. Chromatography 2 used a 30-minute reversed-phase chromatography Acquity UPLC CSH C18 column (1.7 µm, 2.1 mm × 100 mm, 186005297, Waters, Eschborn, Germany) combined with a a VanGuard pre-column (1.7 µm, 2.1 mm × 5 mm, 186005303, Waters), and the gradient was as follows: 0-1 min held at 25% B, 1-3 min from 25% B to 40% B, 3-4 min from 40% B to 50% B, 4-16 min from 50% B to 65% B, 16-17 min from 65% B to 70% B, 17-25 min from 70% B to 75% B, 25-27 min from 75% B to 100% B, and 27-30 min held at 100% B. A 30-minute wash gradient was run between every injection (in parallel with chromatography 1) that used the same gradient as chromatography 1 wash gradient. For both methods mass spectrometer parameters were: source voltage +3250V or -3000 depending on method polarity, sheath gas 40, aux gas 10, sweep gas 1, ion transfer tube temperature 300°C, and vaporizer temperature 275°C. Full scan data were collected using the orbitrap with a scan range of 200-1700 m/z at a resolution of 500,000 and RF lens at 45%. Data dependent MS2 fragmentation was induced in the orbitrap using assisted higher-energy collisional dissociation (HCD) collision energies at 15, 30, 45, 75, and 110% as well as with collision-induced dissociation (CID) at a collision energy of 35%. For both MS2 fragmentations, orbitrap resolution was 15,000 and the isolation window was 1.5 m/z. A m/z 184 mass trigger, indicative of phosphatidylcholines, was used for CID fragmentation. Data dependent MS3 fragmentation was induced in the ion trap with scan rate set at Rapid using CID at a collision energy of 35%. MS3 scans were triggered by specific acyl chain losses for detailed analysis of mono-, di-, and triacylglycerides. Total cycle time was 2 sec. Lipid identifications were assigned using LipidSearch (v5.0, Thermo Fisher Scientific).

For data analysis, peak picking and integration was conducted in Skyline (v22-23) using in-house curated compound lists of accurate mass MS1 and retention time of chemical standards (*9*). For lipidomics studies, the Lipidsearch identifications were used to populate the target compound list for Skyline. For tracing studies, this list was expanded to include all possible carbon isotopologues and natural abundance correction was completed using IsoCorrectR (*11*).

**In vitro acute versus chronic stimulation of T cells**

Chronic stimulation of CD8 T cells was conducted as described (*5*). Briefly, female P14 Thy1.1 C57BL/6 mice aged 8-12 weeks were sacrificed and CD8 (P14) cells were isolated from spleens and peripheral lymph nodes using the EasySep Mouse CD8 T cell isolation kit following manufacturers instructions. Following isolation and flow cytometry-based purity assessment, $5x10^6$ live P14 cells resuspended in TCM containing 10ng/mL IL-2 were placed in overtop chambers of a 6 well dish that were coated 24 hours prior (at 4 °C) with 3 µg/ml anti-mouse CD3 (eBioscience, Clone 145-2C11) and 1 µg/ml anti-mouse CD28 (eBioscience, Clone 37.51) in 1X phosphate-buffered saline (PBS; Wisent, 311-010-CL). After 48h incubation (37 °C incubator, 5% $CO_2$), cells were gently displaced from the coated wells, counted, and $2x10^6$ cells were resuspended in fresh media containing 10 ng/mL IL-2 in a final volume of 5 mL in VIM medium (*8*). For continuous stimulation conditions, cells were placed in 3 wells coated with anti-CD3 and -CD28 antibodies in the same fashion as mentioned above, while for acute stimulation conditions, cells were placed directly in 3 uncoated wells of a 6-well dish in VIM containing IL-2. 48h post incubation, media were changed and cells were seeded in the same fashion as the previous passage.

For flow cytometric assessment of CD8 T cells *in vitro*, $3x10^5$ live cells were centrifuged at 500g for 5 minutes, followed by resuspension in 200 µL TCM with 50 U/mL IL-2. Acute and continuously stimulated cells were seeded in 96 well round bottom plates and re-stimulated with PMA (50 ng/mL) and ionomycin (500 ng/mL) for 4 hours (37°C incubator, 5% $CO_2$), with protein transport inhibitor GolgiStop (1:1500 dilution, BD Biosciences, 5102092KZ) added for the last 2 h, followed by processing for downstream FACS analysis. For unstimulated controls, T cells were only treated with GolgiStop for 2 hours.

For $^{13}C$-tracing experiments, 1-2x$10^6$ T cells were either cultured with for 4h at 37°C in VIM medium containing IL-2 and either 5mM $^{13}C$-glucose/0.85 mM $^{12}C$-βOHB or 5mM $^{12}C$ Glucose/0.85 mM $^{13}C$-βOHB. For unlabeled controls, T cells were cultured in VIM containing 5 mM $^{12}C$-glucose/0.85 mM $^{12}C$-βOHB. Following incubation, cells were pipetted into 15 mL conical tubes and centrifuged at 600g for 3 minutes. Medium was aspirated, then cells washed twice with 5 mL ice cold 0.9% w/v NaCl (saline) and spun at 600g for 2 minutes. Samples were immediately frozen on dry ice for 5 minutes, then transferred to a -80°C freezer until processing for metabolomics.

##### *Ex vivo* Stable Isotope Labeling (SIL) and Metabolomics

*Ex vivo* SIL experiments with CD8$^+$ T cells isolated from mice using liquid chromatography (LC) or gas chromatography (GC) coupled to mass spectrometry (MS) were performed as outlined previously (*1*, *8*). Antigen-specific T cells were isolated from *ad libitum* (AL) or dietary restricted (DR) *Lm*OVA-infected mice (*BDH1*/*OXCT1* DKO or WT mice) using magnetic bead isolation as previously described (*1*, *7*). CD8 T cells were cultured for 2h in VIM medium containing 1.5 mM $^{13}C_4$-βOHB, followed by metabolite extraction as described above. For $^{13}C$ co-tracing experiments, CD8 T cells were cultured for 2h in VIM medium containing 50 U/ml IL-2 and 5 mM $^{13}C_6$-glucose and 1.5 mM 2,4-$^{13}C_2$-βOHB (Cambridge Isotope Laboratories). Identical unlabeled controls were performed using VIM medium containing $^{12}C$-glucose and $^{12}C$-βOHB. After the 2h incubation period, cells were centrifuged at 600g for 4 minutes, media collected and flash-frozen over dry ice, and cell pellets were washed twice with 0.9% NaCl prior to flash-freezing and storage at -80°C for downstream metabolomics analysis.

**Immunoblotting**

Immunoblotting was performed by lysing cells in RIPA buffer containing protease and phosphatase inhibitors (Roche) on ice for a minimum of 30 minutes as previously described (*12*). Protein quantification was done using the Pierce BCA Protein Assay Kit (Thermo Fisher Scientific, Waltham, MA, USA). Equal amounts of protein from whole cell lysates were diluted in Laemmli sample buffer, boiled for 5 minutes, resolved on a 10% SDS-PAGE gel, and transferred to nitrocellulose membranes. Membranes were blocked for 1 hour at room temperature in 5% non-fat milk prepared in TBS-T, followed by overnight incubation at 4°C with primary antibodies against BDH1 or SCOT (1:1000 dilution in 5% non-fat milk). After washing three times in 1x TBS-T (5 minutes per wash), membranes were incubated for 1 hour at room temperature with HRP-conjugated secondary antibodies (diluted in 5% non-fat milk). Membranes were washed again three times with 1x TBS-T before being developed using ECL solution (Cytiva). Details of the antibodies used are provided in the Key Resources Table.

**Bioenergetics (Seahorse) Extracellular Flux assay**

T cell oxygen consumption rate (OCR) and extracellular acidification rate (ECAR) were measured using a Seahorse XF96 Extracellular Flux Analyzer, as previously described (*1*). Antigen specific Thy1.1$^+$ OT-1 CD8$^+$ T cells isolated from *Lm*OVA-infected mice were seeded at 2 x $10^5$ cells per well in Seahorse XF medium containing 5 mM glucose and 0.5 mM glutamine, with or without 1.5 mM βOHB (depending on experimental conditions). Cells were centrifuged onto a poly-D-lysine-coated XF96 plate, and cellular bioenergetics evaluated at 5 minute intervals after the sequential addition of oligomycin (2 μM), FCCP (1.5 μM), rotenone/antimycin A (0.5 μM each), and monensin (20 μM). Data were normalized to cell number. Bioenergetics data interpretation followed the protocols established by Mookerjee *et al.* (*13*).

**Flow cytometry**

Single-cell suspensions were prepared from mouse spleens by mechanical dissociation. Red blood cells (RBCs) were lysed using RBC lysis buffer containing 0.15 M $NH_4Cl$, 10 mM $KHCO_3$, and 0.1 mM EDTA, followed by neutralization with three volumes of TCM. For cell staining, single-cell suspensions were incubated with a cocktail of fluorescently labeled antibodies and dyes as listed in the Key Resources Table. Cell viability was assessed using Fixable Viability Dye eFluor 506 (Thermo Fisher Scientific) according to the manufacturer's protocols. To assess cytokine production following *Lm*OVA infection, splenocytes were harvested from infected mice at 7 dpi were stimulated with 1 μg/mL $OVA_{257\text{-}264}$ peptide (Anaspec) and 50 U/mL interleukin-2 (IL-2) for 4 hours at 37 °C. TIL were restimulated with PMA and ionomycin. GolgiStop protein transport inhibitor (BD Biosciences) was added at a 1:1500 dilution during the last 2 hours of incubation to block cytokine secretion. After stimulation, cells were stained with surface marker antibodies in staining buffer (PBS containing 2% fetal bovine serum [FBS] and 0.02% sodium azide) for 1 hour at 4 °C. Cells were fixed and permeabilized using the Foxp3/Transcription Factor Staining Buffer Set (Thermo Fisher Scientific) at

4°C for 1 hour. Intracellular staining was performed by incubating cells with fluorescently labeled antibodies targeting intracellular markers for either 1h or overnight at 4°C. Flow cytometry data were acquired using a CytoFLEX (Beckman Coulter), Aurora Cytek, or BD Accuri C6 Plus cytometer. Cell sorting was performed on an Astrios (Beckman Coulter) or BD FACSAria Fusion cell sorter. Data analysis was conducted using FlowJo software.

**CITE-Sequencing Experimental Protocol**

C57BL/6J mice aged 10–12 weeks were allocated to either ad libitum (AL) feeding or dietary restriction (DR) for one week prior to tumor cell injection. Following the feeding regimen, each mouse was subcutaneously injected with $5 \times 10^5$ B16-OVA melanoma cells. Thirteen days post-injection, tumors were harvested from the mice for further analysis. To prepare single-cell suspensions, excised tumors were mechanically dissociated and filtered sequentially through 100 µm and then 50 µm cell strainers. Throughout all procedures, cells were kept on ice to preserve viability. The resulting cell suspensions were counted, and cell viability was assessed by trypan blue exclusion. To prevent nonspecific antibody binding, cells were blocked with TruStain FcX™ Plus (BioLegend; Cat# 156603) in Cell Staining Buffer (CSB; BD Biosciences Cat# 420201) for 10 minutes at 4 °C. Each tumor sample was then labeled with a specific antibody (Hashtags 1–8, product information in Key Resource Table) according to the manufacturer's protocol, including appropriate incubation and washing steps.

After hashtag labeling, cells were stained with a panel of fluorescently labeled antibodies targeting surface markers to identify live $CD45^+$ leukocytes while excluding red blood cells (Ter119$^-$), tumor cells (CD105$^-$), and dead cells (DAPI$^+$). This staining facilitated the isolation of the desired cell population during flow cytometric sorting. Cells were sorted by flow cytometry, and live CD45$^+$ cells were collected into tubes containing IMDM supplemented with 10% FBS. Post-sorting, cells were centrifuged and resuspended in CSB, and cell viability reassessed. Based on cell counts, CD45$^+$ cells were adjusted to a concentration of 1 x $10^6$ cells per 50 µL and blocked again with TruStain FcX™ Plus for 10 minutes at 4°C. During this blocking step, TotalSeq™ C antibodies were prepared according to the manufacturer's instructions (BioLegend). The TotalSeq™ C antibodies were then added to the CD45$^+$ cells and incubated for 30 minutes at 4°C to label cell surface proteins for subsequent analysis.

Following antibody staining, cells were washed three times with CSB to remove unbound antibodies. Cells were then resuspended in 1X PBS containing 0.04% pure bovine serum albumin (BSA) at an approximate concentration of 2,200 cells/µL. The prepared cell suspension was used for downstream library preparation and sequencing.

**CITE-Seq Library Preparation**

Libraries were generated and sequenced by the Van Andel Institute Genomics Core. Cells were processed with 10X Chromium Next GEM Single Cell 5' GEM kit v1.1 (10X Genomics, Pleasanton, CA) according to the manufacturer's instructions to target an output of 13,000 cells per sample using a 10X Genomics Chromium Controller. Briefly, single cell suspensions in PBS + 0.04% BSA were assessed for quantity and viability on the CytoFLEX S (Beckman Coulter, Indianapolis, IN), then 20,000 cells per sample were loaded onto the Chromium Controller. Single cells were captured in gel beads in emulsion (GEMs), where they were lysed. Released RNA was barcoded and then converted to cDNA. Using beads, cDNA was separated from the hashtag oligos (HTO) and each were used to generate adapter ligated libraries. Quality and quantity of the finished gene expression and HTO libraries were assessed using a combination of Agilent DNA High Sensitivity chip (Agilent Technologies, Inc.) and QuantiFluor® dsDNA System (Promega Corp., Madison, WI, USA). 2 x 100 bp, paired end sequencing was performed on an Illumina NovaSeq 6000 sequencer (Illumina Inc., San Diego, CA, USA) using an S2, 200 cycle sequencing kit (v1.5) to a minimum depth of 20K reads per cell. Base calling was done by Illumina RTA3 and output was demultiplexed and converted to FastQ format with Cell Ranger (10X Genomics, v3.1.0).

**CITE-Sequencing Quality Control and Processing**

Following library preparation, FastQ files were processed using Cell Ranger version 7.0.1 to map and quantify expression profiles for both the transcriptome and epitopes. Single-cell 5′ paired-end sequencing chemistry was employed, and the sequencing data were aligned to the mouse reference genome m10-2020-A. The resulting raw_feature_bc_matrix, containing both transcript-level and epitope quantifications, was imported into R using the Read10X function from the Seurat package (version 5.0.0). Initial data processing involved separating the gene expression (GEX) data, antibody-derived tags (ADTs), and hashtag oligos (HTOs), which was critical for downstream analysis. A Seurat object was created using the CreateSeuratObject function. Quality control and normalization steps were then performed: HTO data were normalized using the NormalizeData function with the method set to “CLR” (centered log ratio transformation), and the HTODemux function was utilized to classify cells based on HTO signals into “Singlets,” “Doublets,” and “Negative” categories. Further QC filtering was applied by retaining cells with ADT counts of at least 150, RNA counts of at least 200, an HTO classification of “Singlet,” and mitochondrial gene expression percentage less than 20%. The GEX matrix was normalized using the SCTransform function (sctransform version 0.4.1), and ADT data were normalized using the NormalizeData function. Principal component analysis (PCA) was conducted on both RNA and ADT data using the RunPCA function, including all ADTs for dimensionality reduction.

To generate a joint UMAP embedding that integrates both GEX and ADT data, the FindMultiModalNeighbors function was employed, followed by the RunUMAP function, resulting in a weighted nearest neighbor UMAP. Cluster identification was achieved by integrating gene expression profiles, ADT data, and gene expression density visualizations using the plot_density() function from the Nebulosa package (version 1.14.0). Details of cluster identification are provided in **Tables S3–S4** and **Figures S3–S4**. Statistical analyses for violin plots were performed using the Kruskal-Wallis test, followed by pairwise Wilcoxon comparison tests with Bonferroni p-value adjustment. Sequencing data from the single-cell RNA sequencing experiments have been deposited in the Gene Expression Omnibus (GEO) under accession number GSE267070.

**RNA velocity estimation**

To predict single-cell developmental directionality, we performed RNA velocity analysis using the Python implementation of Velocyto, which quantifies ratios of spliced and unspliced transcripts following the method described by La Manno et al (*14*). This approach enabled us to determine the directional changes of cells between dietary conditions. Code used to make RNA velocity plots can be found at https://github.com/rgjcanada/DR_CITESEQ_2025.git.

**Human TIL Single Cell RNA Sequencing**

For analysis of the human scRNA-seq dataset (*15*), count data were normalized and transformed through SCT normalization method in Seurat, with 5000 variable features retained for downstream dimension reduction techniques. Integration of data was performed on the patient level with Canonical Correlation Analysis as the dimension reduction technique. Cells were clustered utilizing the Louvain algorithm with multi-level refinement. Principal component analysis was performed, with the first 50 PCs utilized in UMAP generation. The data was subset to CD8$^+$ T cells, which were identified utilizing the labels provided by Guo *et al.* (*16*) and confirmed via singleR utilizing the ImmGen database (*17*) and cell type annotation with the ProjectTILs T cell atlas (*18*). These CD8$^+$ T cells were subsequently normalized through SCT method, with 3000 variable features retained. Due to the low number of cells on per-patient level, Harmony was utilized to integrate the data at the patient level, rather than Seurat (*19*). PCA and UMAP dimension reduction were performed as above for clusteringinto distinct groups, Teff_ITGB1, Teff_TYROBP, Teff_TRIM34, TCM, Trm_TM, Trm_TMEM2, Trm_BAG3, Trm_ZNF20, IEL, Tex_MKI67, Tex_CLNK, based on markers used by Zhang et al (*20*). To simplify analysis, we combined Teff_ITGB1, Teff_TYROBP, and Teff_TRIM34 into the “T effector (Teff)” cluster, while grouping TCM, Trm_TM, Trm_TMEM2, Trm_BAG3, Trm_ZNF20, IEL into the “T memory

(Tmem)" cluster. Additionally, Tex_MKI67 and Tex_CLNK were combined into the "T exhausted (Tex)" cluster.

**CD8 DAB Staining Quantification**

Full resolution SVS images of HDAB stained tissue sections were imported into QuPath (v0.5.1) (*21*) for analysis. For all tissues to be analyzed, a region of interest (ROI) contouring the tissue was created with the SAM plugin (v0.6.0) for QuPath (*22*, *23*) using the following settings: vit_h (huge) model, foreground with rectangle draw prompt, and single mask output on live mode. To detect nuclei, an average of the deconvolved Hematoxylin, DAB, and Residual channels was generated to create a fluorescence-like single channel image using a custom script (hosted on https://github.com/vaioic). Random regions from the samples were used for CellPose's human-in-the-loop training based on the livecell_cp3 model to improve detection of nuclei containing DAB staining (*24*–*26*). Detections were then generated on the average channel image using the custom trained CellPose-based model using the CellPose plugin for QuPath (v.9.0) (https://github.com/BIOP/qupath-extension-cellpose). The detections and tissue outline were exported as an object geojson and imported to the original HDAB stained image. Detection measurements were generated with QuPath's built in Add Intensity Features with the following settings: pixel size: 0.5 µm; region: ROI; Tile diameter: 0; Channels: Hematoxylin, DAB; Basic features: mean, standard deviation, min & max, median. Nuclei were then classified as positive or negative for DAB staining using the mean DAB signal at a threshold of 0.117. A pixel thresholder was used to measure the area covered by DAB staining. A training image generated from an equal number and size of random rectangles from each image was used for parameter setting. QuPath's Create Thresholder was used with the following settings: Resolution: Very high (1.01 μm/px); Channel: DAB; Prefilter: Gaussian; Smoothing sigma: 1; Threshold: 0.25. The measure option was then run on the tissue outline to quantify the area of the tissue covered by DAB staining.

##### Statistical analysis

Data are presented as mean ± SD for technical replicates or mean ± SEM for biological replicates. Statistical analysis was assessed by GraphPad Prism software (GraphPad) using unpaired Student's t-test or One-Way ANOVA. Statistical significance is indicated in all figures by the following annotations: *P<0.05, **P<0.01, ***P<0.001, ****P<0.0001.

#### KEY RESOURCES TABLE

| REAGENT or RESOURCE | SOURCE | IDENTIFIER |
|---|---|---|
| **Antibodies** | | |
| Hamster monoclonal anti-mouse CD3e (145-2C11) | Thermo Fisher Scientific | Cat# 16-0031-82; RRID: AB_468847 |
| Hamster monoclonal anti-mouse CD28 (37.51) | Thermo Fisher Scientific | Cat# 16-0281-86; RRID: AB_468923 |
| Rat monoclonal anti-mouse CD8a (53-6.7), BUV395 | BD Biosciences | Cat# 563786; RRID: AB_2732919 |
| Rat monoclonal anti-mouse CD8a (53-6.7), BUV737 | BD Biosciences | Cat# 612759; RRID: AB_2870090 |
| Rat monoclonal anti-mouse CD44 (IM7), BUV805 | BD Biosciences | Cat# 741921; RRID: AB_2871234 |

| | | |
|---|---|---|
| Mouse monoclonal anti-mouse NK-1.1 (PK136), Brilliant Violet 605 | BioLegend | Cat# 108739; RRID: AB_2562273 |
| Rat monoclonal anti-mouse CD127 (A7R34), Brilliant Violet 785 | BioLegend | Cat# 135037; RRID: AB_2565269 |
| Rat monoclonal anti-mouse CD3 (17A2), FITC | Thermo Fisher Scientific | Cat# 11-0032-82; RRID: AB_2572431 |
| Rat monoclonal anti-mouse CD4 (RM4-5), FITC | Thermo Fisher Scientific | Cat# 11-0042-82; RRID: AB_464896 |
| Hamster monoclonal anti-mouse KLRG1 (2F1), Alexa Fluor 532 | Thermo Fisher Scientific | Cat# 58-5893-82; RRID: AB_2815282 |
| Rat monoclonal anti-mouse CD19 (1D3), PE | BioLegend | Cat# 152408; RRID: AB_2629817 |
| Mouse monoclonal anti-mouse CD90.1/Thy1.1 (HIS51), PE | Thermo Fisher Scientific | Cat# 12-0900-81; RRID: AB_465773 |
| Mouse monoclonal anti-human/mouse Granzyme B (QA16A02), PE/Dazzle 594 | BioLegend | Cat# 372216; RRID: AB_2728383 |
| Rat monoclonal anti-mouse CD8a (53-6.7), PE-Cyanine7 | Thermo Fisher Scientific | Cat# 25-0081-82; RRID: AB_469584 |
| Rat monoclonal anti-mouse TNF-alpha (MP6-XT22), PE-Cyanine7 | Thermo Fisher Scientific | Cat# 25-7321-82; RRID: AB_11042728 |
| Rat monoclonal anti-mouse CD4 (RM4-5), APC | Thermo Fisher Scientific | Cat# 17-0042-82; RRID: AB_469323 |
| Rat monoclonal anti-mouse IFN-gamma (XMG1.2), APC | Thermo Fisher Scientific | Cat# 17-7311-82; RRID: AB_469504 |
| Rabbit polyclonal anti-β-ACTIN | Cell Signaling Technology | Cat# 4967; RRID: AB_330288 |
| Goat anti-rabbit IgG, HRP-conjugated | Cell Signaling Technology | Cat# 7074; RRID: AB_2099233 |
| BDH1 antibody | Proteintech | Cat# 15417-1-AP, RRID:AB_2274683 |

| | | |
|---|---|---|
| BDH1 antibody | Proteintech | Cat# 67448-1-Ig, RRID:AB_2882682 |
| OXCT1 antibody | Proteintech | Cat# 12175-1-AP, RRID:AB_2157444 |
| Alexa Fluor(R) 700 anti-mouse CX3CR1 | BioLegend | Cat# 149036, RRID:AB_2629606 |
| CD62L (L-Selectin) | BD Biosciences | Cat# 740218, RRID:AB_2739966 |
| Brilliant Violet 711(TM) anti-mouse CD69 | BioLegend | Cat# 104537, RRID:AB_2566120 |
| Pacific Blue(TM) anti-mouse Ly108 | BioLegend | Cat# 134608, RRID:AB_2188093 |
| Brilliant Violet 605(TM) anti-mouse CD279 (PD-1) | BioLegend | Cat# 135219, RRID:AB_1112537 1 |
| CD366 (TIM3) Monoclonal Antibody (RMT3-23), APC, eBioscience | Thermo Fisher Scientific | Cat# 17-5870-82, RRID:AB_2688131 |
| IFN gamma Monoclonal Antibody (XMG1.2), APC, eBioscience | Thermo Fisher Scientific | Cat# 17-7311-82, RRID:AB_469504 |
| TNF alpha Monoclonal Antibody (MP6-XT22), PE-Cyanine7, eBioscience | Thermo Fisher Scientific | Cat# 25-7321-82, RRID:AB_1104272 8 |
| TOX Antibody, anti-human/mouse, PE, REAfinity™ | Miltenyi Biotec | Cat# 130-120-785, RRID:AB_2801785 |
| TCF1/TCF7 (C63D9) Rabbit mAb (Alexa Fluor® 647 Conjugate) | Cell Signaling Technology | Cat# 6709, RRID:AB_2797631 |
| Brilliant Violet 605(TM) anti-T-bet | BioLegend | Cat# 644817, RRID:AB_1121938 8 |
| Goat polyclonal anti-Armenian hamster IgG (H+L), secondary antibody, biotin-conjugated | Thermo Fisher Scientific | Cat# 13-4113-85; RRID: AB_466651 |
| TotalSeq(TM)-CMouse Universal Cocktail, V1.0 | BioLegend | Cat# 199903, RRID:AB_2924498 |
| TotalSeq(TM)-C0301 anti-mouse Hashtag 1 | BioLegend | Cat# 155861<br><br>RRID:AB_2800693 |

| | | |
|---|---|---|
| TotalSeq(TM)-C0302 anti-mouse Hashtag 2 | BioLegend | Cat# 155863, RRID:AB_2800694 |
| TotalSeq(TM)-C0303 anti-mouse Hashtag 3 | BioLegend | Cat# 155865, RRID:AB_2800695 |
| TotalSeq(TM)-C0304 anti-mouse Hashtag 4 | BioLegend | Cat# 155867, RRID:AB_2800696 |
| TotalSeq(TM)-C0305 anti-mouse Hashtag 5 | BioLegend | Cat# 155869, RRID:AB_2800697 |
| TotalSeq(TM)-C0306 anti-mouse Hashtag 6 | BioLegend | Cat# 155871, RRID:AB_2819910 |
| TotalSeq(TM)-C0307 anti-mouse Hashtag 7 | BioLegend | Cat# 155873, RRID:AB_2819911 |
| TotalSeq(TM)-C0308 anti-mouse Hashtag 8 | BioLegend | Cat# 155875, RRID:AB_2819912 |
| TruStain FcX(TM) PLUS (anti-mouse CD16/32) | BioLegend | Cat# 156603, RRID:AB_2783137 |
| Cell Staining Buffer | BioLegend | Cat# 420201 |
| APC anti-mouse CD45 | BioLegend | Cat# 103111, RRID:AB_312976 |
| FITC anti-mouse TER-119/Erythroid Cells | BioLegend | Cat# 116206, RRID:AB_313707 |
| CD105 (Endoglin) Monoclonal Antibody (SN6), PE, eBioscience | Thermo Fisher Scientific | Cat# 12-1057-42, RRID:AB_1311123 |
| InVivoPlus polyclonal Armenian hamster IgG | Bio X Cell | Cat# BP0091, RRID:AB_1107773 |
| InVivoPlus anti-mouse PD-1 (CD279) | Bio X Cell | Cat# BP0033-2, RRID:AB_1107747 |
| **Bacterial and Virus Strains** | | |
| Attenuated (∆actA) *Lm*OVA | John Harty | Haring *et al.*(*27*) |
| **Chemicals, Peptides, and Recombinant Proteins** | | |
| DMEM with 4.5 g/L glucose and L-glutamine, without sodium pyruvate | Wisent Inc. | Cat# 319-015-CL |
| IMDM with L-glutamine & 25 mM HEPES | Wisent Inc. | Cat# 319-105-CL |
| Van Andel Institute-modified IMDM (VIM) | Custom | Kaymak *et al.*(*8*) |

| | | |
|---|---|---|
| Seahorse XF base medium, without phenol red | Agilent Technologies | Part# 103335-100 |
| Nu-Serum IV Culture Supplement | Corning | Cat# 355504; Lot# 2080003 |
| Fetal bovine serum (FBS), heat-inactivated | Corning | Cat# 35-016-CV; Lot# 16821001 |
| Fetal bovine serum (FBS), dialyzed | Corning | Cat# 35-071-CV; Lot# 35071105 |
| Pen Strep (5,000 U/mL penicillin and 5,000 μg/mL streptomycin) | Gibco | Cat# 15070063 |
| 2-mercaptoethanol (55 mM; 1000x) | Gibco | Cat# 21985023 |
| Recombinant murine IL-2 | Peprotech | Cat# 212-12 |
| Recombinant human IL-2 | Peprotech | Cat# 200-02 |
| ImmunoCult Human CD3/CD28 T Cell Activator | StemCell Technologies | Cat# 10971 |
| D-Glucose [U-$^{13}C_6$] | Cambridge Isotopes | Cat# CLM-1396 |
| Sodium D-3-hydroxybutyrate (2,4-$^{13}C_2$) | Cambridge Isotopes | Cat# CLM-3706-1 |
| Sodium D-3-hydroxybutyrate ($^{13}C_4$) | Cambridge Isotopes | Cat# CLM-3853-PK |
| D-Glucose | Sigma-Aldrich | Cat# G8270 |
| L-Glutamine | Sigma-Aldrich | Cat# G3126 |
| Fixable Viability Dye eFluor 506 | Thermo Fisher Scientific | Cat# 65-0866-14 |
| Fixable Viability Dye eFluor 780 | Thermo Fisher Scientific | Cat# 65-0865-14 |
| Violet Proliferation Dye 450 | BD Biosciences | Cat# 562158 |
| BD GolgiStop | BD Biosciences | Cat# 51-2092KZ |
| Foxp3/Transcription Factor Staining Buffer Set | Thermo Fisher Scientific | Cat# 00-5523-00 |
| $OVA_{257-264}$ (SIINFEKL) peptide | AnaSpec | Cat# AS-60193-1 |
| cOmplete, EDTA-free (protease inhibitor cocktail tablets) | Roche | Cat# 11873580001 |

| | | |
|---|---|---|
| PhosSTOP (phosphatase inhibitor cocktail tablets) | Roche | Cat# 4906845001 |
| Monensin sodium salt | Sigma-Aldrich | Cat# M5273 |
| Streptavidin, Alexa Fluor 647 | Thermo Fisher Scientific | Cat# S21374 |
| Phorbol 12-myristate 13-acetate (PMA) | Millipore Sigma | Cat# P1585 |
| Ionomycin calcium salt | Millipore Sigma | Cat# I0634 |
| **Critical Commercial Assays** | | |
| EasySep Mouse T cell Isolation Kit | StemCell Technologies | Cat# 19851 |
| EasySep Mouse CD8$^{+}$ T cell Isolation Kit | StemCell Technologies | Cat# 19853 |
| EasySep Mouse CD90.1 Positive Selection Kit | StemCell Technologies | Cat# 18958 |
| Seahorse XFe96 FluxPak | Agilent Technologies | Part# 102416-100 |
| Seahorse XF Cell Mito Stress Test Kit | Agilent Technologies | Part# 103015-100 |
| Pierce BCA Protein Assay Kit | Thermo Fisher Scientific | Cat# 23225 |
| Pierce Rapid Gold BCA Protein Assay Kit | Thermo Fisher Scientific | Cat# A53227 |
| **Deposited data** | | |
| CITE-Sequencing FASTQ | This manuscript | NCBI GEO: GSE267070 |
| RDS Files (Processed data) | This manuscript | 10.5281/zenodo.13920172 |
| Human Single Cell RNA Sequencing | Zhang L et. al. (*15*) | NCBI GEO: GSE146771 |
| **Experimental Models: Cell Lines** | | |
| 293T cells | ATCC | CRL-3216 |
| MC38-OVA-tdTomato cells | | Luda *et al.*(*1*) |
| B16-OVA cells | | Cordeiro *et al.*(*4*) |

| | | |
|---|---|---|
| EO-771 cells | ATCC | CRL-3461 |
| **Experimental Models: Organisms/Strains** | | |
| C57BL/6J mice | The Jackson Laboratory | RRID: IMSR_JAX:000664 |
| C57BL/6-Tg(TcraTcrb)1100Mjb/J (OT-I) mice | The Jackson Laboratory | RRID: IMSR_JAX:003831 |
| B6.PL-*Thy1*$^{a}$/CyJ (Thy1.1) mice | The Jackson Laboratory | RRID: IMSR_JAX:000406 |
| **B6.Cg-Tg(Cd4-cre)1Cwi/BfluJ (*Cd4-Cre*) mice** | The Jackson Laboratory | RRID: IMSR_JAX:022071 |
| ***Bdh1*$^{fl/fl}$/*Oxct1*$^{fl/fl}$*Cd4-Cre*** | This paper | |
| ***Bdh1*$^{fl/fl}$/*Oxct1*$^{fl/fl}$*Cd4-Cre* OT-I** | This paper | |
| **Oligonucleotides** | | |
| *Bdh1 genotyping primers*:<br>Forward:<br>TGCAGGAATCAGTGCTCTCTCCTAGCA<br>Reverse: GGT GTC AGG GCT GAA GGA TG | This paper | Custom |
| *Oxct1 genotyping primers:*<br>Forward:<br>TATGGGACTCTGGTACAGGAAG<br>Reverse:<br>TGCTGACCGTTAAACTCCCTC | This paper | Custom |
| **Software and Algorithms** | | |
| Adobe Illustrator v28.1 | Adobe | adobe.com/products/illustrator.html |
| Fiji v2.15.1 | | imagej.net/software/fiji |
| FlowJo v9.9.5 | FlowJo LLC | flowjo.com |
| GraphPad Prism v9-10 | GraphPad Software | graphpad.com |
| IncuCyte v2022A Rev1 | Sartorius | sartorius.com/en |
| R v4.4.0 | | cran.r-project.org |
| Python | | V3.12.0 |

**Figure S1**

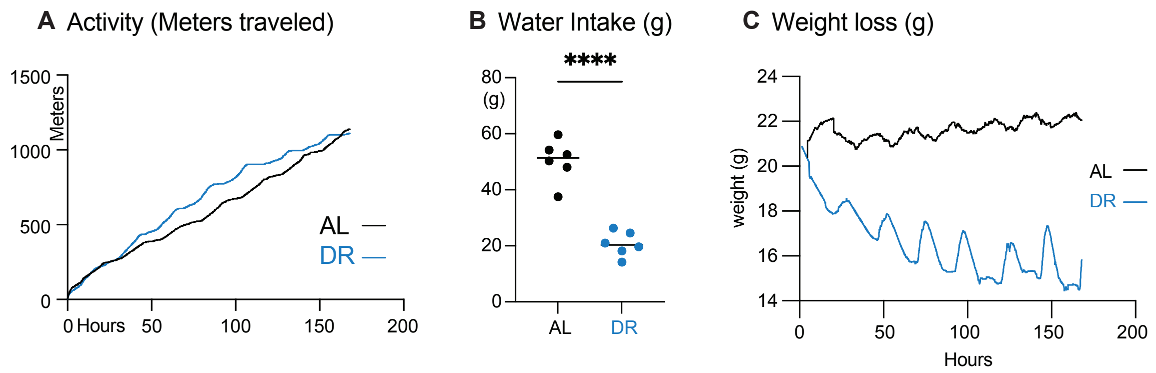


**Figure S1, related to Figure 1. Dietary restriction reduces water intake and body weight without affecting mouse activity levels.**

A) Total distance traveled (meters) by mice on ad libitum (AL) or dietary restriction (DR) feeding regimens. Activity was measured using metabolic Promethion cages over 168 hours (n = 6 mice per group).

B) Total water intake (grams) over one week for mice on AL or DR feeding regimens. Data represent the mean ± SEM (n = 6 mice per group).

C) Change in body weight (grams) over time (hours) for mice on AL or DR conditions. Data represent the mean ± SEM (n = 6 mice per group).

*P<0.05, **P<0.01, ***P<0.001, ****P<0.0001.

#### Figure S2

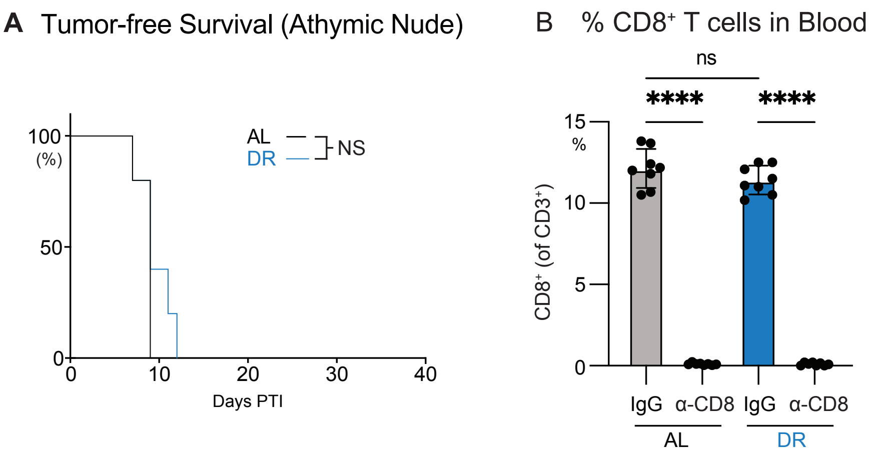


**Figure S2, related to Figure 1. Dietary restriction does not delay tumor onset in immunodeficient mice.**

A) Kaplan-Meier plot comparing tumor onset (tumor volume ≥250 mm³) in B16-OVA melanoma-bearing athymic nude mice fed ad libitum (AL) or dietary restriction (DR) diets. Statistical significance was assessed by log-rank test (n = 5 mice per group).

B) Bar plot showing serum immunoglobulin levels in mice administered anti-IgG or anti-PD1 antibodies under AL or DR conditions. Data represent the mean ± SEM (n = 8 mice per group).

*P<0.05, **P<0.01, ***P<0.001, ****P<0.0001.

### Figure S3

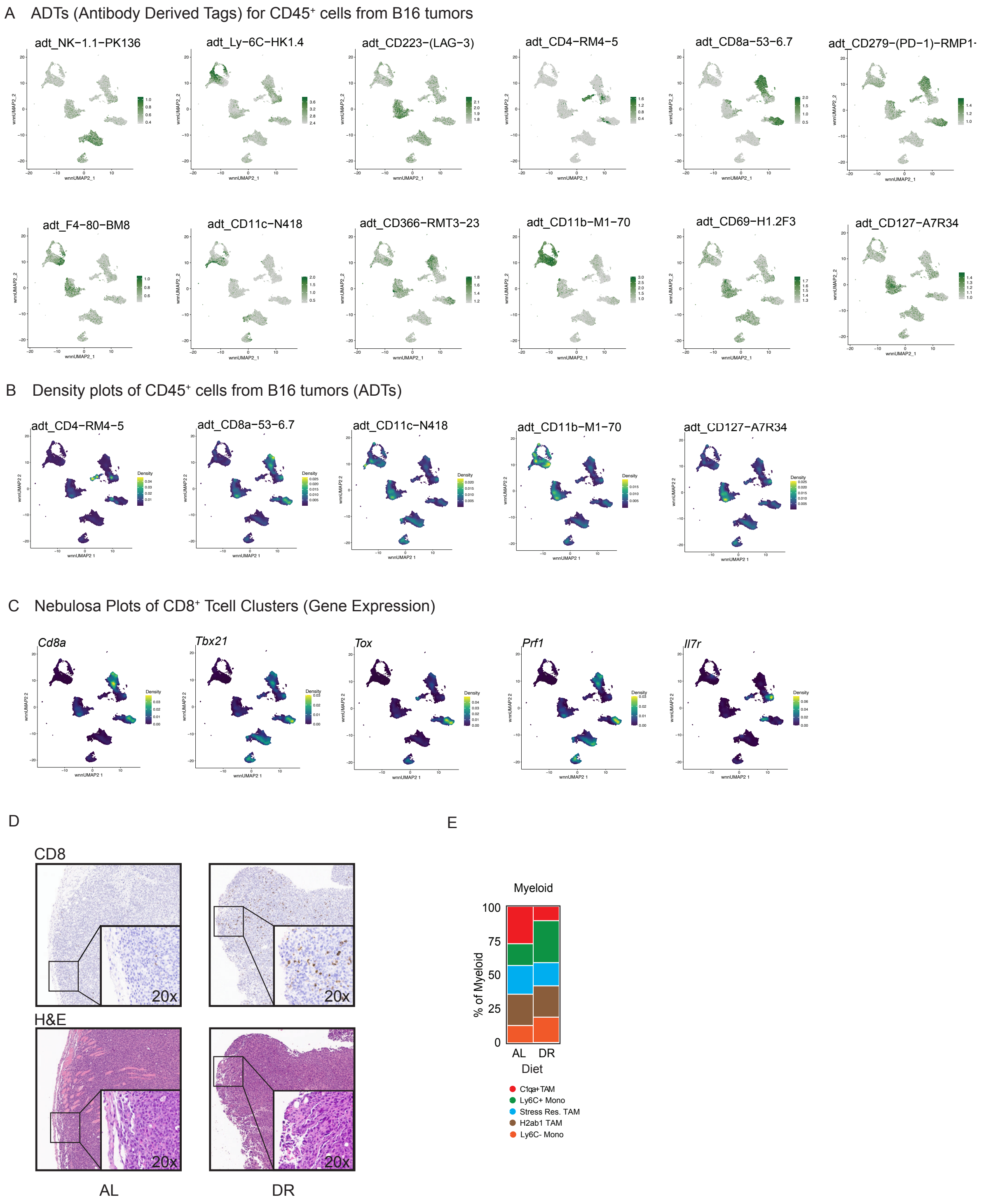


**Figure S3, related to Figure 1. Dietary restriction enhances CD8$^{+}$ T cell infiltration and alters immune cell composition in B16 melanoma tumors.**

A) Weighted nearest neighbor Uniform Manifold Approximation and Projection (wnnUMAP) of 45,455 CD45$^+$ tumor-infiltrating cells from B16 melanoma tumors of mice on AL or DR diets (wnnUMAP combined; n = 4 mice per diet). Each plot represents the expression of key antibody-derived tags (ADTs) for markers including NK1.1, Ly6C, LAG3 (CD223), CD4, CD8, PD1, F4/80, CD11c, TIM3 (CD366), CD11b, CD69, and CD127.

B) Density plots generated using the Nebulosa package, showing the expression density of ADTs (CD4, CD8, CD11c, CD11b, CD127) among activated CD8$^+$ T cells from B16 tumors (combined AL and DR conditions).

C) Density plots of gene expression for *Cd8a*, *Tbx21*, *Tox*, *Prf1*, and *Il7r* among activated CD8$^+$ T cells from B16 tumors (joint plot of AL and DR conditions).

D) Histological analysis of B16 tumors from AL- or DR-fed mice 14 days post-tumor implantation. Immunohistochemical staining for CD8$^+$ T cells (brown) and hematoxylin and eosin (H&E) staining of representative tumor sections are shown. Insets are at 20X magnification; original images are at 4X magnification.

E) Stacked bar graph showing the percentage breakdown of myeloid cell populations in B16 tumors from AL and DR mice.

*P<0.05, **P<0.01, ***P<0.001, ****P<0.0001.

### Figure S4

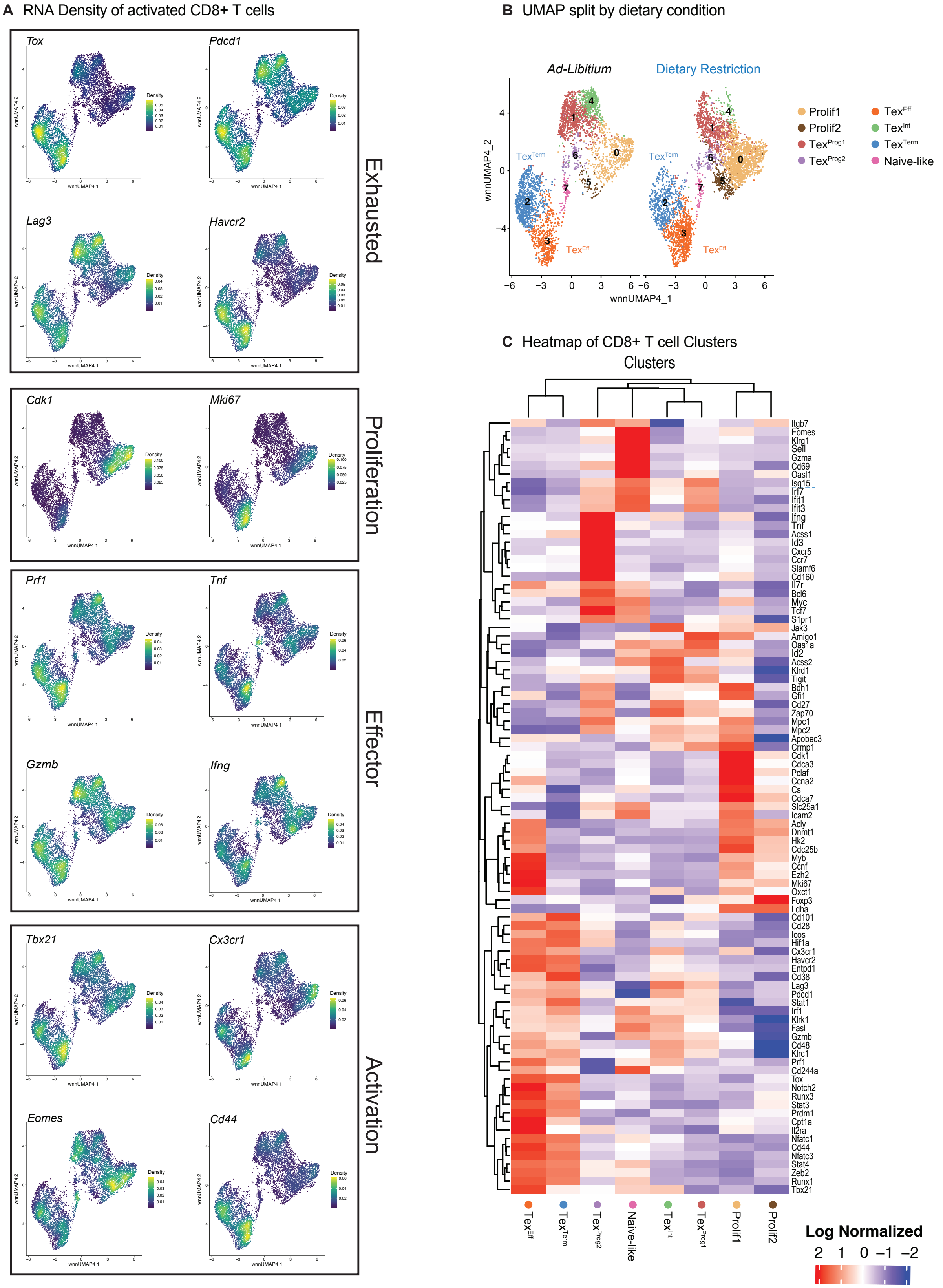
A RNA Density of activated CD8+ T cells
Tox
Pdcd1
Lag3
Havcr2
Exhausted
Cdk1
Mki67
Proliferation
Prf1
Tnf
Gzmb
Ifng
Effector
Tbx21
Cx3cr1
Eomes
Cd44
Activation
Density
wnnUMAP4 1
wnnUMAP4 2
B UMAP split by dietary condition
Ad-Libitium
Dietary Restriction
wnnUMAP4_2
wnnUMAP4_1
Prolif1
Prolif2
TexProg1
TexProg2
TexEff
TexInt
TexTerm
Naive-like
C Heatmap of CD8+ T cell Clusters
Clusters
Itgb7
Eomes
Klrg1
Sell
Gzma
Cd69
Oasl1
Isg15
Irf7
Ifit1
Ifit3
Ifng
Tnf
Acss1
Id3
Cxcr5
Ccr7
Slamf6
Cd160
Il7r
Bcl6
Myc
Tcf7
S1pr1
Jak3
Amigo1
Oas1a
Id2
Acss2
Klrd1
Tigit
Bdh1
Gfi1
Cd27
Zap70
Mpc1
Mpc2
Apobec3
Crmp1
Cdk1
Cdca3
Pclaf
Ccna2
Cs
Cdca7
Slc25a1
Icam2
Acly
Dnmt1
Hk2
Cdc25b
Myb
Ccnf
Ezh2
Mki67
Oxct1
Foxp3
Ldha
Cd101
Cd28
Icos
Hif1a
Cx3cr1
Havcr2
Entpd1
Cd38
Lag3
Pdcd1
Stat1
Irf1
Klrk1
Fasl
Gzmb
Cd48
Klrc1
Prf1
Cd244a
Tox
Notch2
Runx3
Stat3
Prdm1
Cpt1a
Il2ra
Nfatc1
Cd44
Nfatc3
Stat4
Zeb2
Runx1
Tbx21
TexEff
TexTerm
TexProg2
Naive-like
TexInt
TexProg1
Prolif1
Prolif2
Log Normalized
2 1 0 −1 −2

**Figure S4, related to Figure 2. Dietary restriction alters CD8$^+$ T cell clusters and gene expression profiles in the tumor microenvironment.**

A) RNA density plots of activated CD8$^+$ T cell clusters represented on a wnnUMAP, displaying the density of genes associated with exhaustion, proliferation, effector function, and activation gene sets (joint plot of AL and DR conditions).

B) wnnUMAP of activated CD8$^+$ tumor-infiltrating lymphocytes (TILs) divided by dietary conditions (AL: 3,348 cells; DR: 3,657 cells). Prominent CD8$^+$ T cell clusters are indicated (n = 4 mice per group).

C) Heatmap showing $Log_2$-normalized expression of key genes highlighting population differences among cell clusters in TILs isolated from B16 tumors (combined AL and DR conditions).

#### Figure S5

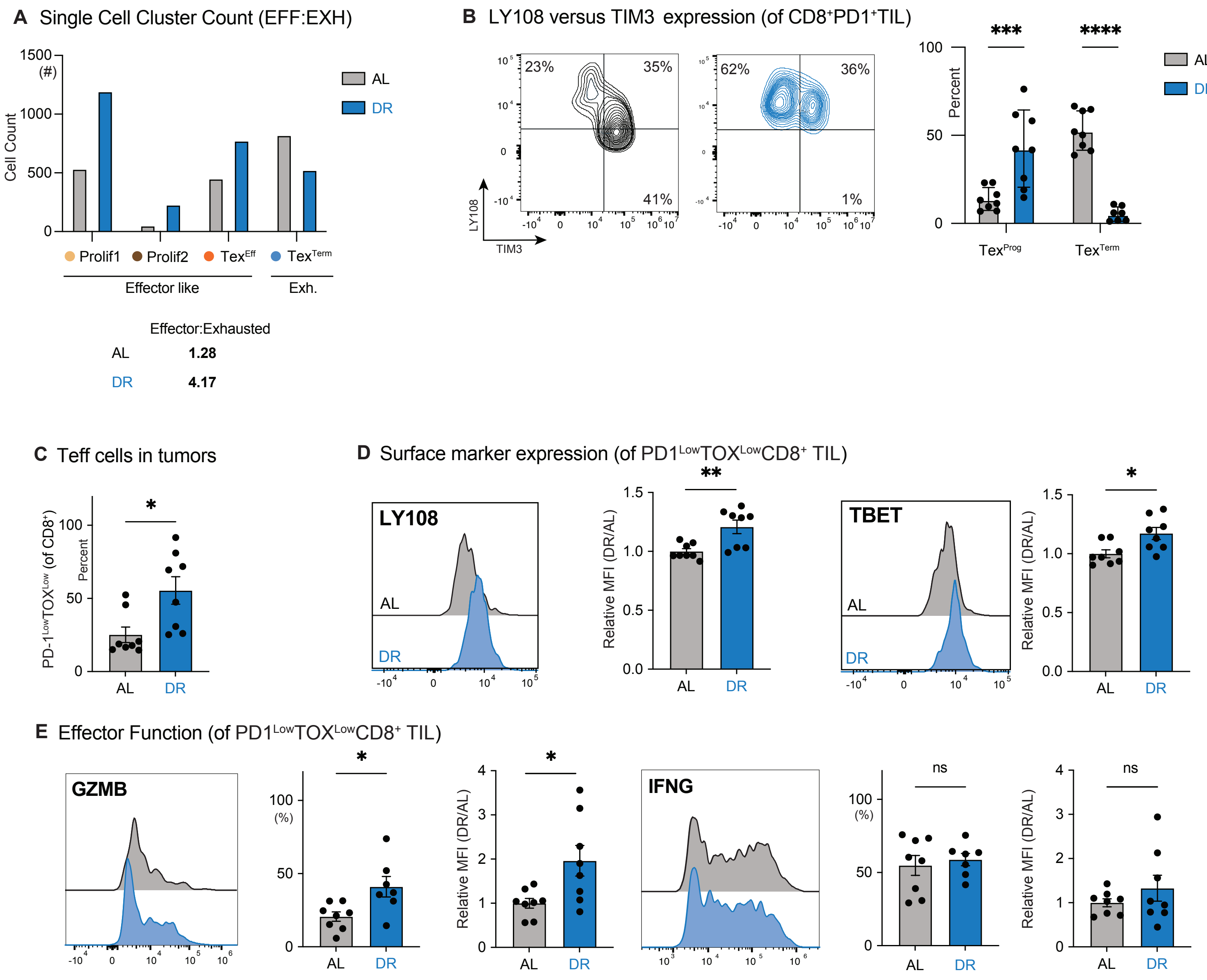


**Figure S5, related to Figure 2. Dietary restriction shifts CD8⁺ T cell populations towards effector-like cells and reduces terminal exhaustion.**

A) Bar plot depicting cell counts from CITE-sequencing experiments for Prolif1, Prolif2, TexEff, and TexTerm CD8⁺ TIL clusters under AL and DR conditions. The Effector:Exhausted ratio is displayed based on cell counts.

B) Expression of LY108 versus TIM3 among $PD1^+CD8^+$ TILs from B16 tumors. *Left,* representative flow cytometry plots showing LY108 and TIM3 expression. *Right*, bar graph showing percentages of progenitor ($Tex^{Prog}$, $LY108^+TIM3^-$) and terminal ($Tex^{Term}$, $LY108^-TIM3^+$) Tex cell subsets under AL and DR conditions. Data represent the mean ± SEM (n = 8 mice per group).

C) Bar graph showing the percentage of $PD1^{Low}TOX^{Low}$ $CD8^+$ T cells among TILs from B16 tumors under AL and DR conditions. Data represent the mean ± SEM (n = 8 mice per group).

D) Representative histograms of LY108 and TBET expression in $PD1^{Low}TOX^{Low}$ $CD8^+$ TILs from B16 tumors. Bar plots show relative geometric mean fluorescence intensity (gMFI) compared to AL controls. Data represent the mean ± SEM (n = 8 mice per group).

E) Representative histograms of Granzyme B (GZMB) and IFN-γ expression in $PD1^{Low}TOX^{Low}$ $CD8^+$ TILs from B16 tumors. Bar plots show relative gMFI compared to AL controls. Data represent the mean ± SEM (n = 8 mice per group).

*P<0.05, **P<0.01, ***P<0.001, ****P<0.0001.

**Figure S6**

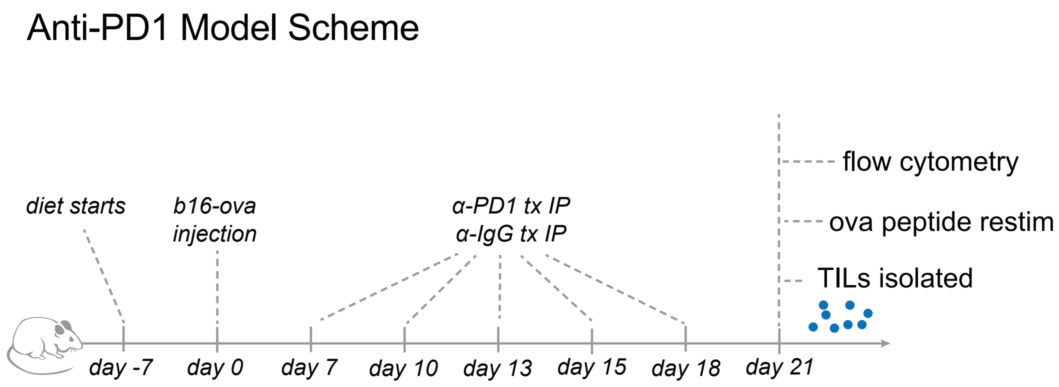


**Figure S6, related to Figure 2. Experimental design for anti-PD1 treatment in mice under dietary restriction.**

Schematic representation of anti-IgG and anti-PD1 antibody treatment in mice bearing B16 melanoma tumors under ad libitum (AL) or dietary restriction (DR) feeding regimens. Antibody treatments (200 μg/dose) were administered every three days for a total of five injections, starting on day 7 post-tumor implantation.

#### Figure S7

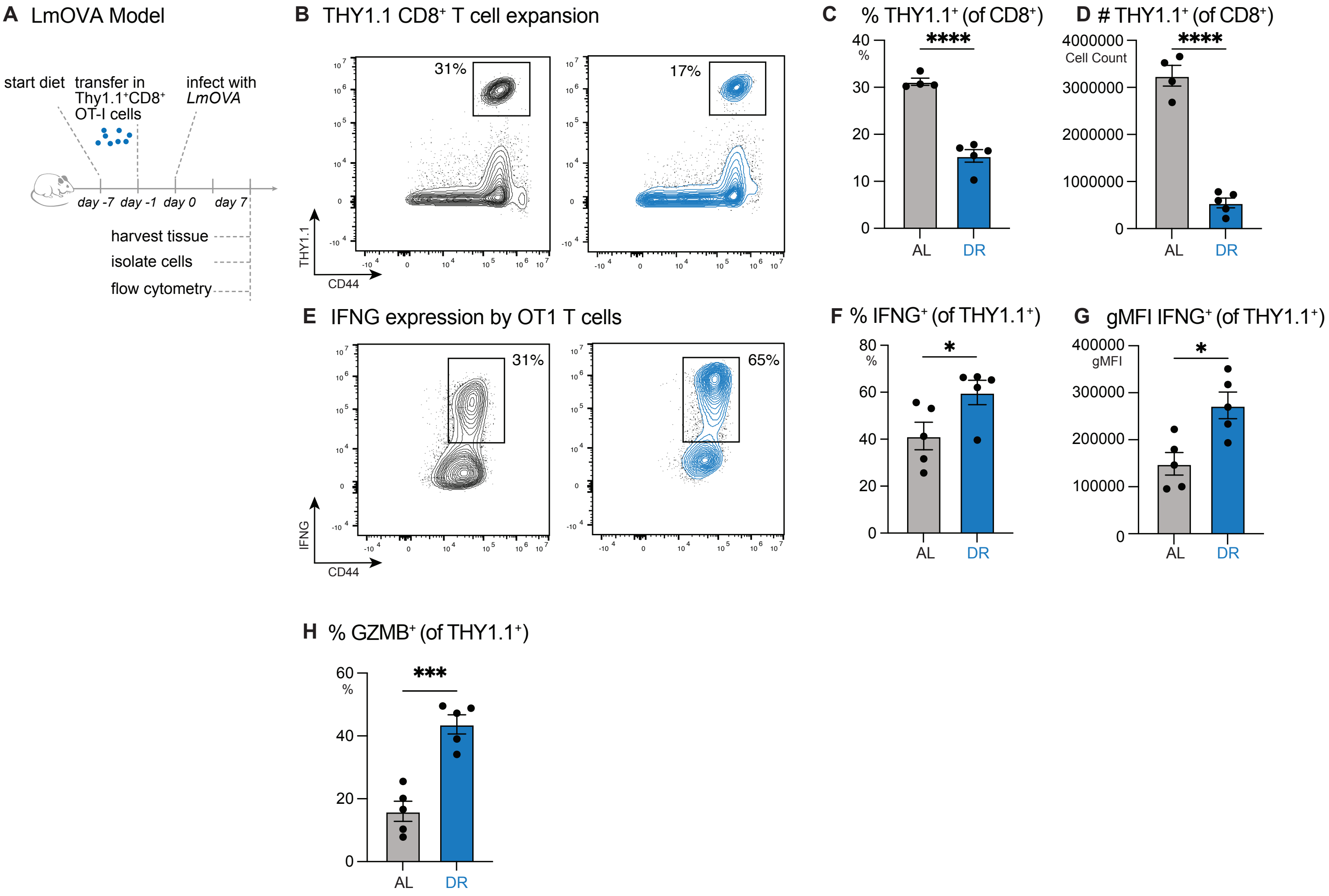


**Figure S7, related to Figure 3. Dietary restriction enhances expansion and effector function of antigen-specific CD8⁺ T cells in response to LmOVA infection.** A) Schematic depicting the experimental setup of adoptive OT-I T cell transfer coupled with attenuated *Lm*OVA infection. Mice were started on AL or DR diets six days before OT-I adoptive transfer. On day 0, mice were injected with LmOVA via tail vein. Seven days post-infection (dpi), spleens were harvested for downstream analyses, including effector function, cell expansion, and $^{13}C$ metabolic tracing.

B) Representative flow cytometry plots showing expansion of splenic OT-I $CD8^+$ T cells (THY1.1 versus CD44) 7 dpi.

C-D) Bar plots showing the percentage (C) and absolute number (D) of $THY1.1^+$ antigen-specific $CD8^+$ T cells expanded under AL or DR conditions at 7 dpi. Data represent the mean ± SEM (n = 4–5 mice per group).

E) Representative flow cytometry plots showing IFN-γ production and CD44 expression in splenic OT-I $CD8^+$ T cells ($CD8^+THY1.1^+$) under AL or DR conditions at 7 dpi.

F-G) Bar plots showing (F) the percentage of IFN-$\gamma^+$ cells and (G) geometric mean fluorescence intensity (gMFI) of IFN-γ in $CD8^+THY1.1^+$ T cells under AL or DR conditions at 7 dpi. Data represent the mean ± SEM (n = 5 mice per group).

H) Bar plot showing the percentage of Granzyme $B^+$ cells among $CD8^+THY1.1^+$ T cells under AL and DR conditions at 7 dpi. Data represent the mean ± SEM (n = 5 mice per group).

*P<0.05, **P<0.01, ***P<0.001, ****P<0.0001.

### Figure S8

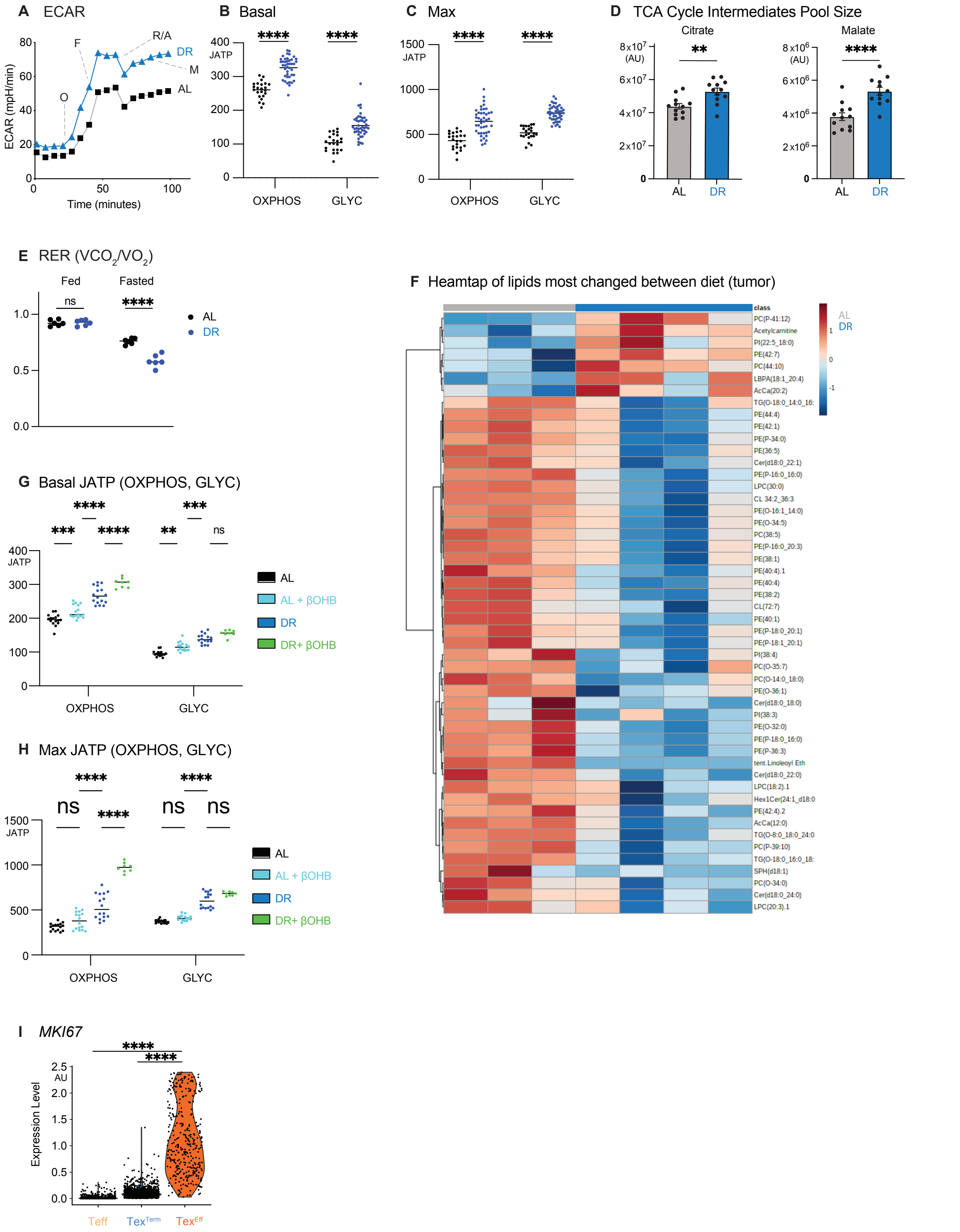

A ECAR
ECAR (mpH/min)
Time (minutes)
O
F
R/A
M
DR
AL
B Basal
C Max
JATP
OXPHOS
GLYC
****
D TCA Cycle Intermediates Pool Size
Citrate
Malate
(AU)
**
****
AL
DR
E RER (VCO2/VO2)
Fed
Fasted
ns
F Heamtap of lipids most changed between diet (tumor)
class
PC(P-41:12)
Acetylcarnitine
PI(22:5_18:0)
PE(42:7)
PC(44:10)
LBPA(18:1_20:4)
AcCa(20:2)
TG(O-18:0_14:0_16:
PE(44:4)
PE(42:1)
PE(P-34:0)
PE(36:5)
Cer(d18:0_22:1)
PE(P-16:0_16:0)
LPC(30:0)
CL 34:2_36:3
PE(O-16:1_14:0)
PE(O-34:5)
PC(38:5)
PE(P-16:0_20:3)
PE(38:1)
PE(40:4).1
PE(40:4)
PE(38:2)
CL(72:7)
PE(40:1)
PE(P-18:0_20:1)
PE(P-18:1_20:1)
PI(38:4)
PC(O-35:7)
PC(O-14:0_18:0)
PE(O-36:1)
Cer(d18:0_18:0)
PI(38:3)
PE(O-32:0)
PE(P-18:0_16:0)
PE(P-36:3)
tent.Linoleoyl Eth
Cer(d18:0_22:0)
LPC(18:2).1
Hex1Cer(24:1_d18:0
PE(42:4).2
AcCa(12:0)
TG(O-8:0_18:0_24:0
PC(P-39:10)
TG(O-18:0_16:0_18:
SPH(d18:1)
PC(O-34:0)
Cer(d18:0_24:0)
LPC(20:3).1
G Basal JATP (OXPHOS, GLYC)
***
**
AL + βOHB
DR+ βOHB
H Max JATP (OXPHOS, GLYC)
I MKI67
Expression Level
AU
Teff
Tex^Term
Tex^Eff

**Figure S8, related to Figure 3. Dietary restriction enhances metabolic activity and mitochondrial function in $CD8^+$ T cells.**

A) Extracellular acidification rate (ECAR) plot of antigen-specific $CD8^+THY1.1^+$ T cells isolated from LmOVA-infected mice (7 dpi) under AL or DR conditions. Data represent the mean ± SD (n = 24–46 technical replicates).

B) Basal ATP production rates (J_ATP) from oxidative phosphorylation (OXPHOS) and glycolysis (GLY) for $CD8^+THY1.1^+$ T cells under AL or DR conditions. Data represent the mean ± SD (n = 24–46 technical replicates).

C) Maximal ATP production rates (J_ATP) from OXPHOS and GLY for $CD8^+THY1.1^+$ T cells under AL or DR conditions. Data represent the mean ± SD (n = 24–46 technical replicates).

D) Bar plots showing the pool sizes of TCA cycle intermediates citrate and malate in antigen-specific $CD8^+THY1.1^+$ T cells from LmOVA-infected mice (7 dpi) under AL or DR conditions. Data represent the mean ± SEM (n = 11–12 mice per group).

E) Mean respiratory exchange ratio (RER; $VCO_2/VO_2$) of mice on AL or DR feeding regimens measured using Promethion metabolic cages. “Fed” represents the average peak RER post-feeding, while “Fasted” represents the lowest RER during fasting periods. Data represent the mean ± SEM (n = 6 mice per group).

F) Heatmap showing the most significant lipid changes ($Log_2$ fold change) in tumors isolated from mice on AL or DR diets 14 days post tumor implantation (n = 3–4 mice per group).

G) Theoretical basal ATP production rates (J_ATP) from OXPHOS and GLY in $CD8^+$ T cells cultured with or without β-hydroxybutyrate (βOHB) under AL or DR conditions. Data represent the mean ± SEM (n = 8–16 technical replicates).

H) Theoretical maximal ATP production rates (J_ATP) from OXPHOS and GLY under the same conditions as in (G). Data represent the mean ± SEM (n = 8–16 technical replicates).

*P<0.05, **P<0.01, ***P<0.001, ****P<0.0001.

**Figure S9**

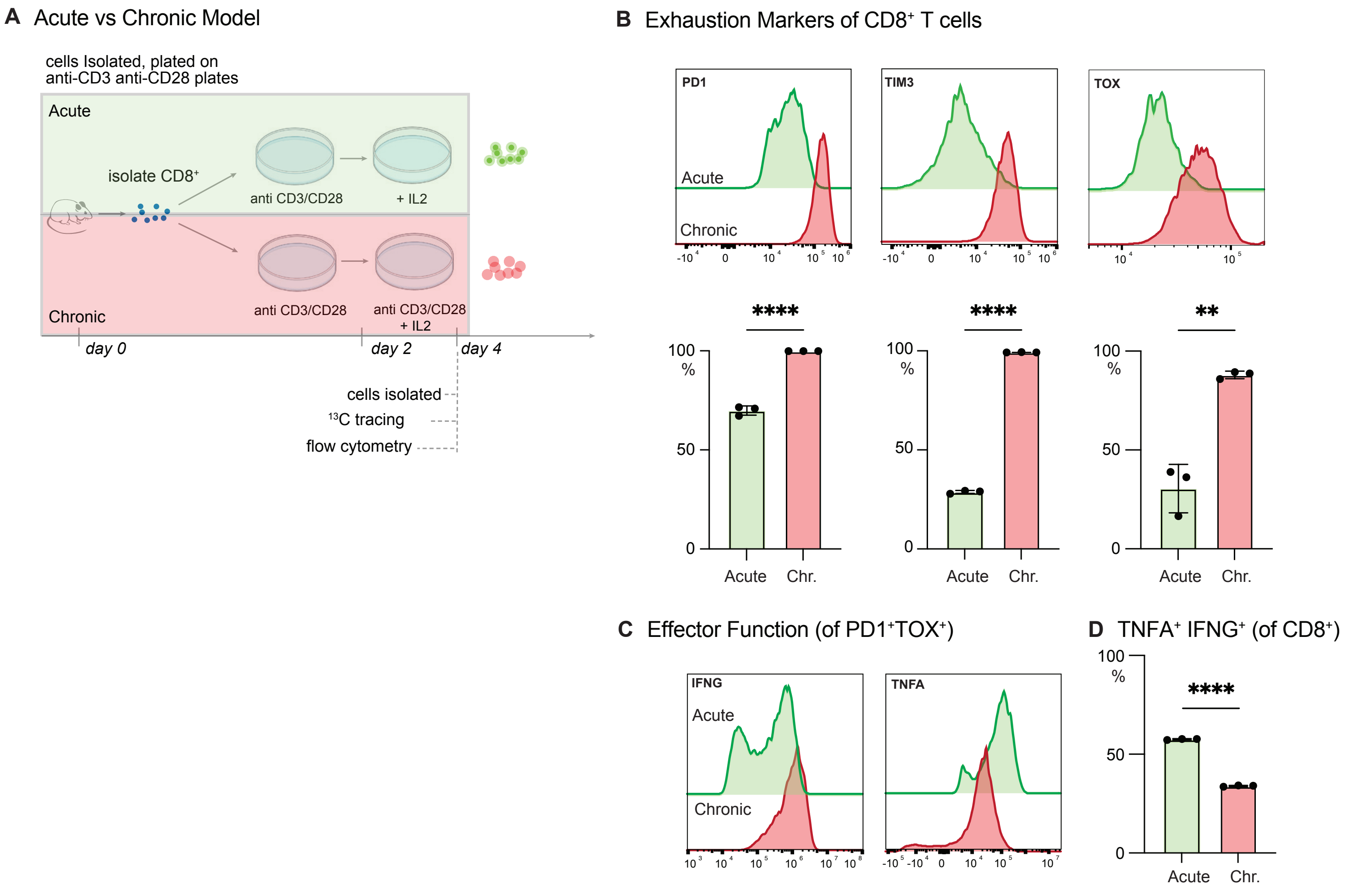


**Figure S9, related to Figure 4. Chronic antigen stimulation induces T cell dysfunction in vitro.**

A) Schematic illustrating the acute versus chronic in vitro stimulation assay, adapted from (*5*). CD8$^+$ T cells isolated from C57BL/6 mice were activated with anti-CD3 and anti-CD28 antibodies (1 µg/mL and 3 µg/mL, respectively) for two days. “Acute” cells were removed from the activation plate and cultured with IL-2, while “chronic” cells were replated on fresh anti-CD3 and anti-CD28-coated plates for an additional 2–4 days.

Cells were then assessed for markers of T cell exhaustion, activation, and effector function.

B) Inhibitory receptor expression on in vitro-stimulated T cells. *Top*, representative histograms of exhaustion marker expression (PD1, TIM3, TOX) in $CD8^+$ T cells under acute and chronic stimulation conditions. *Bottom*, bar graphs of expression levels for each exhaustion marker. Data represent the mean ± SEM (n = 3 mice per group).

C) Representative histograms of effector cytokine production (IFN-γ and TNF-α) in $CD8^+$ T cells under acute and chronic stimulation conditions.

D) Bar plot showing the percentage of TNF-$\alpha^+$IFN-$\gamma^+$ double-positive $CD8^+$ T cells under acute and chronic stimulation conditions. Data represent the mean ± SEM (n = 3 mice per group).

*P<0.05, **P<0.01, ***P<0.001, ****P<0.0001.

### Figure S10

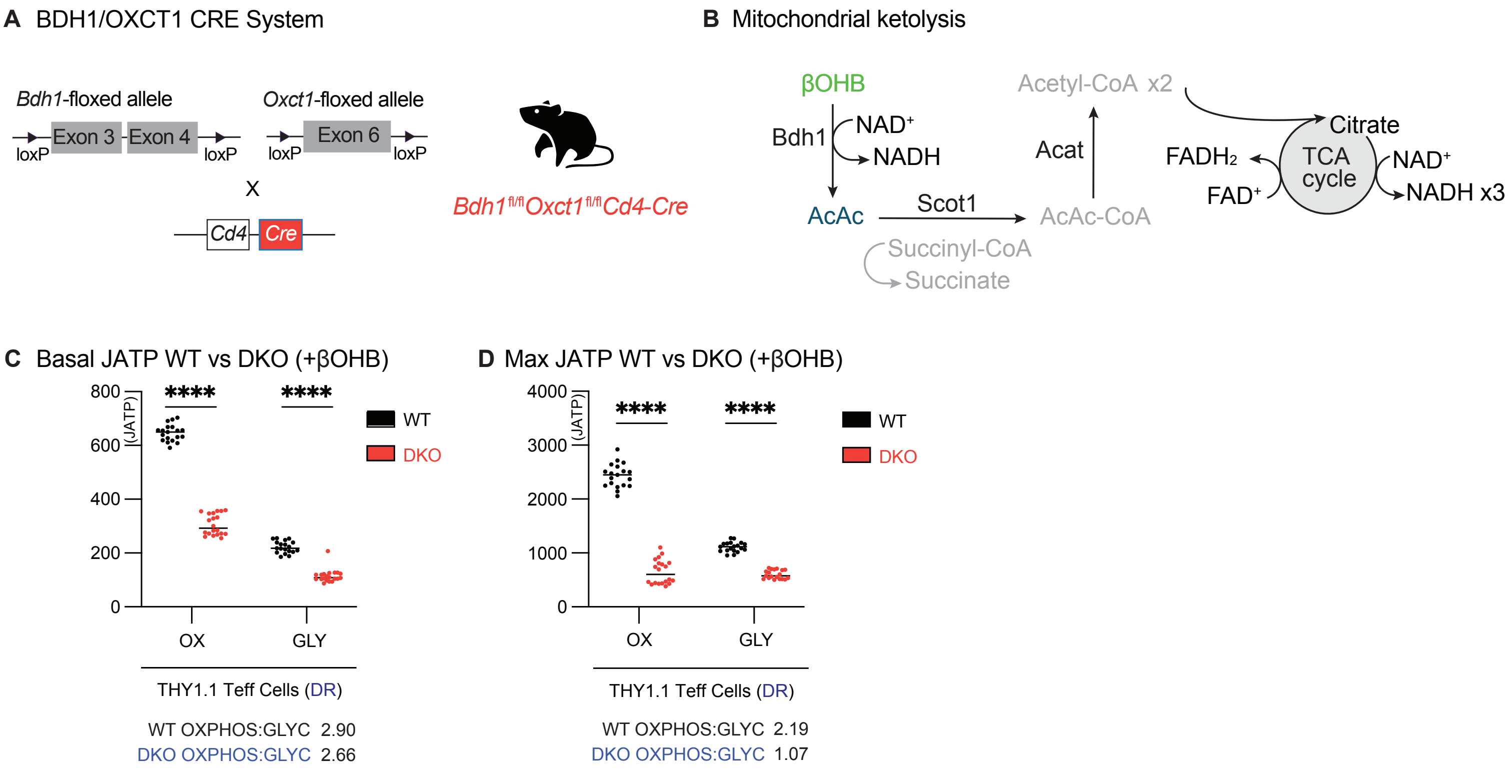


**Figure S10, related to Figure 4. Ketolysis-deficient T cells exhibit impaired metabolic function and fail to respond to dietary restriction.**

A) Schematic detailing the generation of *Bdh1*$^{fl/fl}$*Oxct1*$^{fl/fl}$ *Cd4*-Cre mice. Bdh1-floxed mice were crossed with Oxct1-floxed mice to generate double-floxed mice expressing Cre recombinase under the *Cd4* promoter, resulting in T cells unable to metabolize ketone bodies.

B) Diagram illustrating mitochondrial ketolysis pathways. β-Hydroxybutyrate (βOHB) is converted to acetoacetate (AcAc) by BDH1. AcAc is then converted to acetoacetyl-CoA by SCOT1 (*Oxct1*), ultimately leading to acetyl-CoA production for entry into the TCA cycle and ATP generation.

C) Basal ATP production rates (J_ATP) from oxidative phosphorylation (OXPHOS) and glycolysis (GLY) in antigen-specific $CD8^{+}THY1.1^{+}$ T cells from wild-type (WT) and double knockout (DKO; *Bdh1*$^{fl/fl}$*Oxct1*$^{fl/fl}$ Cd4-Cre) mice under DR conditions. T cells were isolated from LmOVA-infected mice at 7 dpi. Data represent the mean ± SEM (n = 19–20 technical replicates).

D) Maximal ATP production rates (J_ATP) from OXPHOS and GLY under the same conditions as in (C). Data represent the mean ± SEM (n = 19–20 technical replicates). *P<0.05, **P<0.01, ***P<0.001, ****P<0.0001.
